# Improved SILAC quantification with data independent acquisition to investigate bortezomib-induced protein degradation

**DOI:** 10.1101/2020.11.23.394304

**Authors:** Lindsay K Pino, Josue Baeza, Richard Lauman, Birgit Schilling, Benjamin A Garcia

**Affiliations:** Department of Biochemistry and Biophysics, University of Pennsylvania, Philadelphia, PA, U.S.A; Buck Institute for Research on Aging, Novato, CA, U.S.A

**Keywords:** protein turnover, pulse SILAC, protein degradation, data independent acquisition, quantitative proteomics

## Abstract

Stable isotope labeling by amino acids in cell culture (SILAC) coupled to data-dependent acquisition (DDA) is a common approach to quantitative proteomics with the desirable benefit of reducing batch effects during sample processing and data acquisition. More recently, using data-independent acquisition (DIA/SWATH) to systematically measure peptides has gained popularity for its comprehensiveness, reproducibility, and accuracy of quantification. The complementary advantages of these two techniques logically suggests combining them. Here, we develop a SILAC-DIA-MS workflow using free, open-source software. We determine empirically that using DIA achieves similar peptide detection numbers as DDA and that DIA improves the quantitative accuracy and precision of SILAC by an order of magnitude. Finally, we apply SILAC-DIA-MS to determine protein turnover rates of cells treated with bortezomib, a 26S proteasome inhibitor FDA-approved for multiple myeloma and mantle cell lymphoma. We observe that SILAC-DIA produces more sensitive protein turnover models. Of the proteins determined differentially degraded by both acquisition methods, we find known ubiquitin-proteasome degrands such as HNRNPK, EIF3A, and IF4A1/EIF4A-1, and a slower turnover for CATD, a protein implicated in invasive breast cancer. With improved quantification from DIA, we anticipate this workflow making SILAC-based experiments like protein turnover more sensitive.

## INTRODUCTION

There are generally two approaches to quantitative bottom-up proteomics: those involving isotopic labeling and those label-free (1,2). The former includes strategies such as isobaric labeling using chemical tags and metabolic labeling, such as stable isotope labeling by amino acids in cell culture (SILAC), in which proteins are metabolically labeled in vivo with heavy isotope-containing amino acids, pooled together prior to sample preparation (3). Quantitative SILAC workflows have been typically analyzed by data dependent acquisition (DDA) using MS1 precursor ion abundances for quantification. Here we present a novel strategy using data independent acquisition (DIA) and MS2 chromatogram quantification, where SILAC samples are prepared individually and acquired by DIA separately (4). Then, the two individual MS2 chromatograms are compared to determine differential abundance. An advantage of using metabolic isotopic labeling is the ability to multiplex the samples, and the mass spectrometry acquisition of pooled samples reduces technical variation. However, a clear advantage of DIA is that the MS2-level quantification is typically more reproducible and accurate (5).

Interest in multiplexing DIA experiments by isotopic labeling has led to the exploration of neutron-encoded (NeuCode) SILAC (NeuCoDIA) (6) and mass defect based four-plex data-independent acquisition (MdFDIA) (7). Isobaric tagging (e.g. TMT, iTRAQ) is not possible in DIA acquisition modes because reporter ions from multiple precursors become convoluted and cannot be linked back to the precursor of origin (8). While precursor mass-shifting multiplexing approaches seem promising, there is still a need for fundamental benchmarking to explicitly compare SILAC-DDA versus SILAC-DIA.

Benchmarking is especially necessary because, in theory, SILAC increases the complexity of a sample by the number of labels used. A conventional two-plex SILAC experiment with a light and a heavy proteome combined together is then twice as complex as the equivalent label-free experiment. Due to this increased complexity, there is an expectation that conventional SILAC-DIA/SWATH approaches should underperform conventional SILAC-DDA workflows (9). While comparisons of SILAC-DDA versus DIA-MShave been explored (10) and SILAC labeling is supported in DIA analysis softwares such as Spectronaut (11), there are few explicit comparisons of SILAC-DDA versus SILAC-DIA. However, it has been shown that SILAC increases the reproducibility DIA quantifications (12), and there are experimental designs that require the incorporation of metabolic labels like SILAC including Isotopic Differentiation of Interactions as Random or Targeted (I-DIRT) for protein-protein interactions (13) and pulse SILAC for protein turnover.

Pulse SILAC (pSILAC) is the predominant method to study protein turnover at the proteome scale, enabling the discrimination of preexisting and newly synthesized proteins (14–16). Using pSILAC, the cell’s translational machinery is used to incorporate stable isotope labeled amino acids present in the cell media, thereby marking newly synthesized proteins. Over time, as proteins are synthesized and degraded (also called protein turnover), the levels of the newly synthesized proteins increase while the preexisting proteins decline allowing protein half lives to be determined. This approach has been used to explore proteostasis in a number of biological systems, including differentiation (17), cancer homeostasis (18), drug treatment response (19–21), and the accumulation of “old” proteins in aging (22).

Given the broad range of biological systems that benefit from pulse SILAC turnover studies, and considering the quantitative benefits of DIA-MS approaches, we sought to accelerate turnover studies by validating the hyphenation of these two technologies and establishing a freely available analysis workflow.

## EXPERIMENTAL SECTION

### Cell culture medium

Cell culture medium was prepared from Dulbecco’s Modified Eagle Media (DMEM) (Thermo Fisher) supplemented with 10% dialyzed fetal bovine serum (FBS) (Atlanta Biologicals) and 1% penicillin & streptomycin (Gibco, Life Technologies Corporation, Grand Island, NY, USA). For SILAC experiments, DMEM for SILAC (Thermo Fisher) which is deficient in arginine, and lysine is supplemented with 10% dialyzed fetal bovine serum (dFBS) and 1% Pen/Strep. Light SILAC media is supplemented with L-arginine HCl (84 mg/L), L-lysine HCl (146 mg/L) and L-proline (1000 mg/L). Heavy SILAC media is supplemented with with L-arginine-^13^C_6_, ^15^N_4_ HCl (88.2 mg/L), L-lysine-^13^C_6_, ^15^N_2_ HCl (190.59 mg/L) and L-proline (1000 mg/L) (Cambridge Isotope Laboratories, Andover, MA). Media components were mixed and sterile-filtered through a 0.22 um PES membrane filter (Millipore).

### HeLa SILAC labeling for benchmarks

HeLa cells were cultured in heavy- and light-SILAC culture media continuously for 11 days. The labeled media were removed and the cells were quickly washed twice with sterile PBS at room temperature, scraped off the dishes, pelleted by centrifugation, snap-frozen in liquid nitrogen, and stored at −80 °C.

### Bortezomib pulse-SILAC labeling

Primary human foreskin fibroblasts (HFF) were cultured in DMEM, 10% FBS, 1% Pen/Strep. Prior to pulse-SILAC labeling, HFF cells were conditioned to the SILAC media formulation over four cell passages. With each passage, HFF cells were cultured with an increasing amount of SILAC media which increased in increments of 25%. Passage 1: 75% DMEM, 25% SILAC DMEM; passage 2: 50% DMEM, 50% SILAC DMEM; passage 3: 25% DMEM, 75% SILAC DMEM; passage 4: 100% SILAC DMEM. After culture in light media, HFF were supplemented with DMSO or 1000 pM Bortezomib and concurrently the media was switched to SILAC DMEM. Cells were harvested together after 0, 2, 4, 8, 10, 24, 48, 72, 168, and 500 hours of labeling, washed in PBS, and stored in −80 °C until further analysis.

### Human cell line protein preparation

Cell pellets were resuspended in lysis buffer (8M urea, 75mM NaCl, 50mM Tris pH 8, 1mM EDTA pH 8) and sonicated 3x for 30s, resting on ice in between. Protein concentration was estimated by BCA (Pierce BCA Protein Assay Kit, Thermo Scientific, Rockford, IL, USA). Denatured proteins were reduced with 5 mM DTT, alkylated with 15 mM IAA, and digested overnight with 1:50 trypsin (Promega). Peptides were desalted using an MCX protocol (Oasis MCX cartridge 1cc/30 mg LP, Waters Corporation) and dried down by speed-vac. To generate light/heavy mixtures, light and heavy HeLa peptides were reconstituted to 1 ug/ul and the concentrations adjusted to 1:1 using TIC. Peptides were used to construct a calibration curve spanning three orders of magnitude via five serial dilutions (**Supplementary Table 1**).

### E. coli protein preparation

*E. coli* pellets were available that were either previously labeled with stable isotope labeled lysine residue (^13^C_6_-^15^N_2_-Lys) growing them in heavy SILAC media from Cambridge Isotopes (heavy samples), or they were grown in regular media (light samples). We then processed isolated frozen bacterial pellets. Cell pellets of the heavy and light *E. coli* strains were suspended in 6 mL of PBS and centrifuged at 4°C, 15,000 g for 20 min. The firm cell pellet was collected and re-suspended and denatured in a final solution of 6 M urea, 100 mM Tris, 75 mM NaCl.

Protein lysates containing 1 mg of protein were reduced with 20 mM DTT in 100 mM Tris (37°C for 1 h), and subsequently alkylated with 40 mM iodoacetamide in 100 mM Tris (30 min at RT in the dark) (Sigma Aldrich, St. Louis, MO). Samples were diluted 10-fold with 100 mM Tris (pH 8.0) and incubated overnight at 37°C with sequencing grade trypsin (Promega, Madison, WI) added at a 1:50 enzyme:substrate ratio (wt/wt). Subsequently, samples were acidified with formic acid and desalted using HLB Oasis SPE cartridges (Waters, Milford, MA) (23). Proteolytic peptides were eluted, concentrated to near dryness by vacuum centrifugation, and re-suspended chromatographic (aqueous) buffer A. To generate light/heavy mixtures, light and heavy *E. coli* peptides were diluted spanning 400x range (**Supplementary Table 2**).

### High pH reversed phase (HPRP) fractionation

Off-line peptide fractionation was performed using a 10-step gradient of acetonitrile in ammonium formate, as follows. Columns packed with C-18 (Micro SpinColumn, Harvard Apparatus Cat# 74-4601 (Holliston, MA, USA)) were first activated with methanol, conditioned with 80% acetonitrile in 0.1% formic acid, and equilibrated with 0.1% trifluoroacetic acid. Tryptic peptides (750 ug) from the equimolar proteome mix (50% light HeLa, 50% heavy HeLa) were acidified with trifluoroacetic acid and loaded onto the C-18 column. The peptides were eluted in 10 fractions: 10%, 12.5%, 15%, 17.7%, 20%, 22.5%, 25%, 30%, 40%, and 80% acetonitrile in 100mM ammonium formate pH 10. Each fraction was desalted using an MCX protocol (Oasis MCX cartridge 1cc/30 mg LP, Waters Corporation (Milford, MA, USA)) and dried down by speed-vac.

### Orbitrap liquid chromatography-mass spectrometry for human cell line samples

Peptides were analyzed with a Thermo Dionex UPLC coupled with a Thermo Q-Exactive HFX tandem mass spectrometer. We used an in-house pulled column created from 75 μm inner diameter fused silica capillary packed with 3 μm ReproSil-Pur C18 beads (Dr. Maisch) to 30◻cm. Solvent A was 0.1% formic acid in water, while solvent B was 0.1% formic acid in 80% acetonitrile. For each injection, we loaded approximately 1◻μg peptides and separated them using a 90-minute gradient from 5 to 35% B, followed by a 50◻min washing gradient.

For data dependent acquisition (DDA) analysis, a Top20 method was used (default charge state 3, minimum AGC target 5e4, charge exclusion 1 and >8, and dynamic exclusion 15s) with full MS (resolution 120,000; AGC 1e6, maximum IT 40 ms) and data dependent-MS2 (resolution 15,000; AGC 2e5, max IT 40 ms, isolation window 2.0 m/z, NCE 27)).

For data independent acquisition (DIA) analysis of the dilution series and bortezomib experiments, we performed chromatogram library experiments as described in Searle *et al*. Briefly, we acquired 6 chromatogram library acquisitions with 4◻m/z DIA spectra (4◻m/z precursor isolation windows at 30,000 resolution, AGC target 1e6, maximum inject time 60◻ms, 27 NCE) using a staggered (also referred to as overlapping) window pattern from narrow mass ranges using window placements optimized by Skyline (i.e., 398.43–502.48, 498.48–602.52, 598.52–702.57, 698.57–802.61, 798.61–902.66, and 898.6–1002.70◻m/z). We acquired corresponding precursor spectra matching the range (i.e., 390–510, 490–610, 590–710, 690–810, 790–910, and 890–1010 m/z) using an AGC target of 1e6 and a maximum inject time of 60 ms were interspersed every 25 MS/MS spectra.

For all single-injection acquisitions, the Thermo Q-Exactive HFX was configured to acquire either 75◻×◻8◻m/z (covering 400-1,000 m/z) precursor isolation window DIA spectra (15,000 resolution, AGC target 1e6, maximum inject time 20◻ms, 27 NCE) using an optimized staggered window pattern. Precursor spectra (target range ± 15 m/z at 60,000 resolution, AGC target 1e6, maximum inject time 60◻ms) were interspersed every 75 MS/MS spectra. Isolation window schemes for the pulse SILAC experiments have been previously described (24) and the additional windowing schemes benchmarked in this work including method settings are detailed in **Supplemental Table 3**.

### QqTOF Liquid-Chromatography-Mass Spectrometry Acquisitions of E. coli samples

Briefly, samples were analyzed by reverse-phase HPLC-ESI-MS/MS using an Eksigent Ultra Plus nano-LC 2D HPLC system (Dublin, CA) with a cHiPLC system (Eksigent) which was directly connected to a quadrupole time-of-flight (QqTOF) TripleTOF 6600 mass spectrometer (SCIEX, Concord, CAN). After injection, peptide mixtures were loaded onto a C18 pre-column chip (200 μm x 0.4 mm ChromXP C18-CL chip, 3 μm, 120 Å, SCIEX) and washed at 2 μl/min for 10 min with the loading solvent (H2O/0.1% formic acid) for desalting. Subsequently, peptides were transferred to the 75 μm x 15 cm ChromXP C18-CL chip, 3 μm, 120 Å, (SCIEX), and eluted at a flow rate of 300 nL/min with a 3 h gradient using aqueous and acetonitrile solvent buffers (Burdick & Jackson, Muskegon, MI).

For quantification, all *E. coli* peptide samples were analyzed by data-independent acquisition (DIA), using 64 variable-width isolation windows (5,25). The variable window width is adjusted according to the complexity of the typical MS1 ion current observed within a certain m/z range using a DIA ‘variable window method’ algorithm (more narrow windows were chosen in ‘busy’ m/z ranges, wide windows in m/z ranges with few eluting precursor ions). DIA acquisitions produce complex MS/MS spectra, which are a composite of all the analytes within each selected Q1 m/z window. The DIA cycle time of 3.2 sec included a 250 msec precursor ion scan followed by 45 msec accumulation time for each of the 64 variable SWATH segments.

### DIA data analysis

Mass spectrometry data files were demultiplexed and converted to MZML using MSConvert (version 3.0.18) (26). For single-shot, “direct” DIA experiments, MZML files were searched against the human reference proteome (20350 entries, accessed 2019/10/16) using Walnut (version 0.9.5). For pulseSILAC experiments (dilution series, bortezomib), the processing workflow is depicted schematically in **Figure 1**. First, narrow window GPF-DIA-MS data was searched against a predicted spectral library to generate a chromatogram library (27,28). The chromatogram library was used to search the endogenous, light SILAC peptides in each of the curve points using EncyclopeDIA (version 0.9.0). The resulting elib file was imported into Skyline as a library, the detected light peptides used to populate the Target List, and heavy SILAC pairs were associated with each detected peptide sequence. Demultiplexed MZML data files were then imported into Skyline to extract light and heavy SILAC chromatograms. For a full tutorial on analyzing SILAC-DIA with this method, see **Supplementary Note 1**.

**Figure 1.**
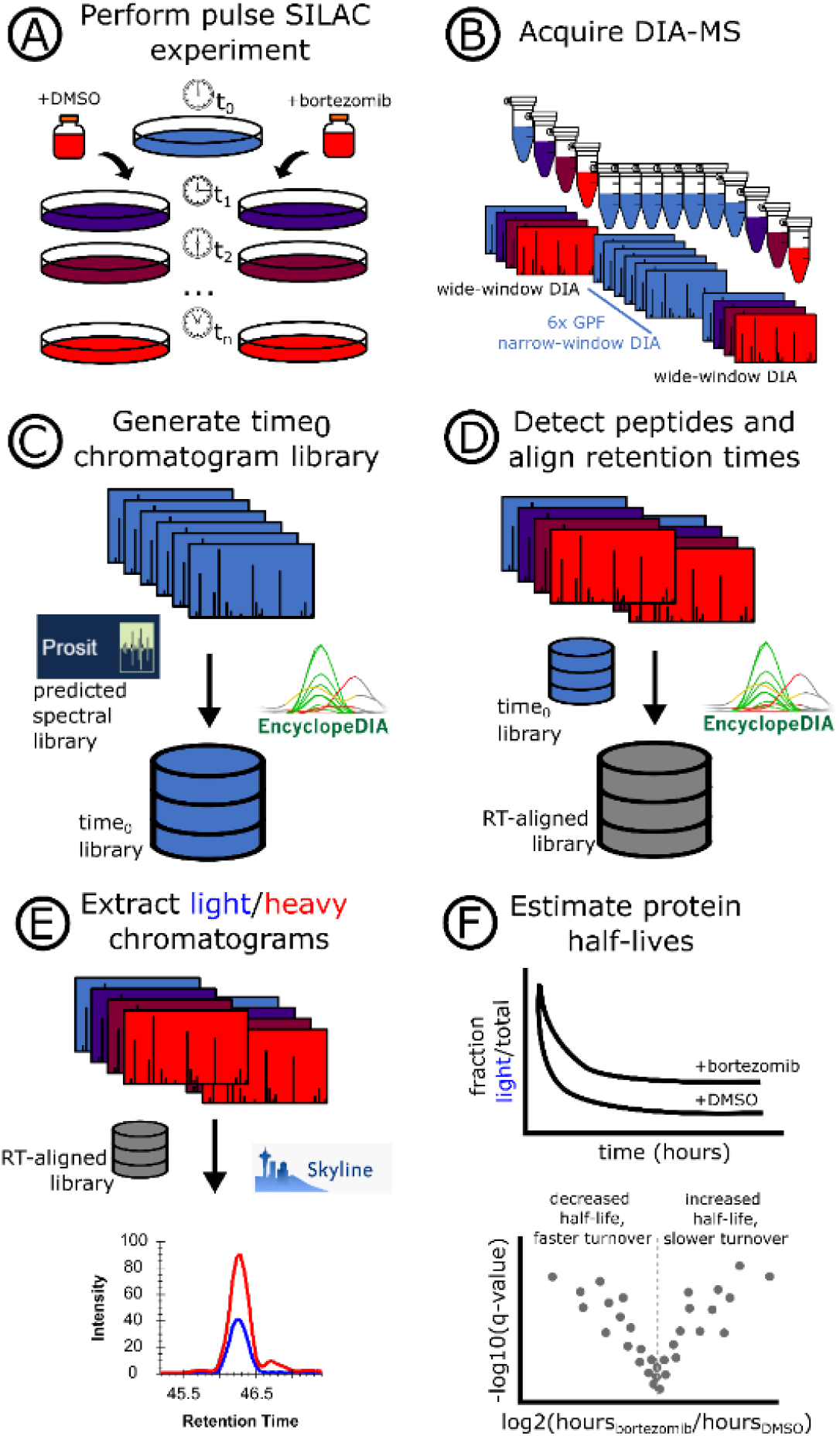
An approach for quantifying pulse SILAC peptides using free, open-source software. A pulse SILAC experiment is performed **(A)** and the data is acquired following a chromatogram library approach **(B)** in which the time_0_ sample(s) are injected multiple times with gas phase fractionation (GPF). The GPF time_0_ spectra files are searched against a predicted spectral library using EncyclopeDIA, creating a time_0_ chromatogram library **(C)** which is then used to align the light peptide retention times across the entire pulse SILAC experiment **(D)**. Using the database of retention time-aligned light peptide detections, the heavy peptides are paired and chromatograms extracted using Skyline **(E)**. Quantitative values for light and heavy SILAC peptides are used to fit protein turnover models and assess statistical significance **(F)**.

### DDA data analysis

RAW files were converted to MZML using MSConvert (version 3.0.18). MetaMorpheus (version 0.0.310) (29) was run with the following parameters: trypsin, 2 max missed cleavages, minimum peptide length 7; fixed carbamidomethyl on C and U, variable oxidation on M; precursor mass tolerance 5 ppm, product mass tolerance 20 ppm; HCD fragmentation. SILAC/SILAM quantification (multiplex) was selected using R(+10.008) & K(+8.014) labels, with “quantify unlabeled peptides/proteins” enabled.

### Data availability

The RAW files, converted MZML files, Metamorpheus output files, Encyclopedia elib files, and Skyline documents have been deposited in ProteomeXChange Consortium (30) via the Panorama (31) partner repository with the identifiers PXD022659 (ProteomeXchange) and https://panoramaweb.org/silac-dia.url (Panorama).

### Statistical determination of turnover rate

Ratios were calculated each for each replicate. Specifically, for each replicate, we use the Skyline-reported Total Area Fragment value (MS2, y-ions only) for Light and Heavy isotopes to calculate the ratio. For the labeling methods in this work, it was crucial to only use y-ions because only that fragmentation series includes the N-terminal SILAC label, ensuring that the fragment m/z is specific to the light or heavy precursor. Protein turnover was determined by fitting the peptide-level data for each protein to an exponential function by nonlinear least squares (“nls” function in the R package “stats”, version 4.0.2) as described by Welle *et al* (32) to obtain the first-order degradation rate constant (*k*_*deg*_) for each protein. Specifically,

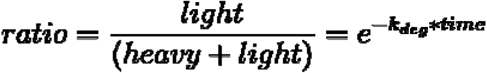

in which *time* is the duration of culture in the heavy media. Each protein turnover model was allowed to run for a maximum of 1000 iterations with all peptide-level data reported for each protein group, else *k*_*deg*_ was reported as missing. The result is a *k*_*deg*_ value for each peptide with light and heavy quantifications in each condition (here, DMSO and bortezomib treatments).

The *k*_*deg*_ is then converted to the time to protein degradation (*halflife*_*hours*_), as estimated by each peptide and in each condition, by

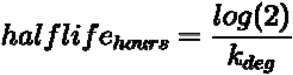

The half lives (above) were then fit to a linear model to determine significantly differential half-lives under bortezomib treatment.

The code is available on Github (https://github.com/lindsaypino/silac-dia.git).

## RESULTS AND DISCUSSION

Despite interest in multiplexing DIA, such as by SILAC-DIA, there is concern about splitting the isotopic distributions of paired precursor species. To illustrate the theory, we simulated isotopic envelopes from the human proteome **(Supplemental Figure 1)**. In an ideal scenario, the heavy isotopic envelope will still fall entirely within the DIA isolation window adjacent to its endogenous paired species. However, there are also undesirable scenarios where one of the SILAC pairs has an isotopic envelope split across two adjacent DIA windows. The concern then, which has been raised by others including Ludwig *et al* 2018 (9), is whether these split isotopic envelopes will affect DIA peptide detections or quantification.

We first approached addressing this theoretical concern by assessing the question of detection in SILAC-DDA vs SILAC-DIA experiments. To do so, we constructed three benchmark samples: a 100% SILAC light proteome, a 100% SILAC heavy proteome, and a 1:1 equimolar mix of SILAC light and heavy. We acquired these three benchmark samples in triplicate by both DDA and DIA, then analyzed the samples using MetaMorpheus and Walnut, respectively. We found that the number of detections in the DDA and the DIA samples is comparable **(Figure 2A)**. The number of detections made by DDA in the 50/50% sample is much lower than in either 100% sample, likely owing to the increased complexity in the 50/50% sample. By doubling the complexity of the sample, roughly 50% of the precursor ions presented to a DDA method will be the redundant SILAC pairs, pointing to a potential pitfall of using DDA for SILAC applications. With this in mind, in single-shot samples, it does not appear that isotopic envelope splitting negatively impacts the number of detections by DIA.

**Figure 2.**
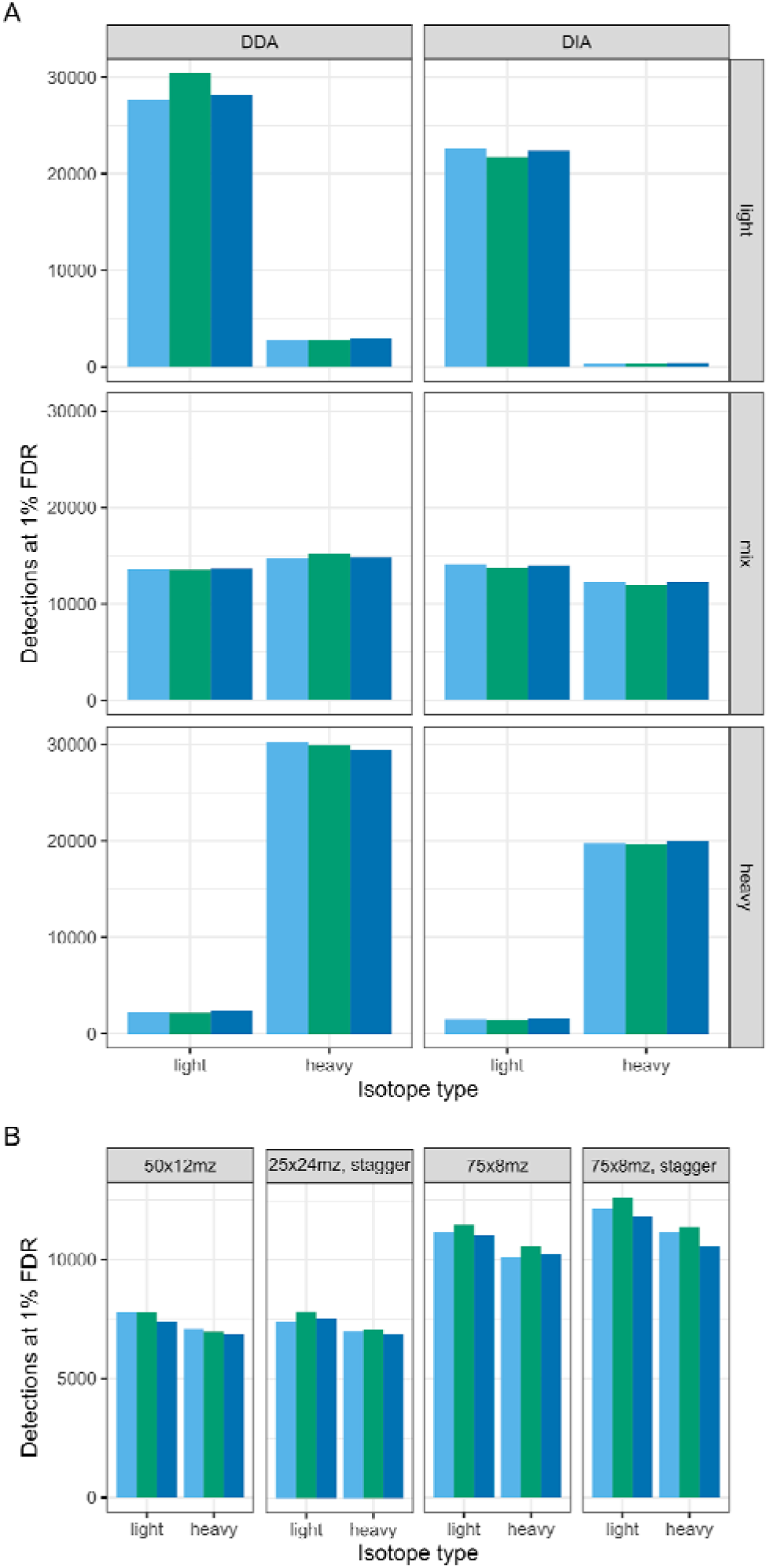
Comparison of PSM and peptide detections in SILAC-DDA vs SILAC-DIA. (A) Detections in three SILAC proteome samples (100% light SILAC, 100% heavy SILAC, and 50/50% light/heavy SILAC “mix”) are compared from three replicates each of DDA and DIA (windowing scheme of 75×8 m/z, staggered). For DDA analysis, the unique PSMs at 1% FDR are used, without using match-between-runs across samples; for DIA analysis, the unique peptides at 1% FDR are used. (B) The number of unique peptides detected at 1% FDR in each technical triplicate of four DIA windowing schemes is compared for the same 50/50% light/heavy SILAC sample. Schemes are described by the number of isolation windows (e.g. “25”) followed by the width of the isolation windows (e.g. “x24mz”), and whether the isolation windows are staggered (“stagger”) or fixed.

We then tested four DIA windowing schemes to determine the effect of isolation window width and staggering on peptide detections. We hypothesized that, if isotopic envelope splitting across windows affected peptide detection, then wider, fixed windows would improve detections. However, we find that narrow and staggered windows achieve better detections than wider, fixed windows **(Figure 2B)**, further supporting that isotopic envelope splitting across DIA isolation windows does not greatly affect peptide detections in empirical data.

Next, we determined whether SILAC would affect quantitative accuracy. For these experiments, we used similar benchmark samples as above (e.g. a 100% light- and a 100% heavy-labeled HeLa proteome), but rather than combining them 1:1, we made serial dilutions spanning several orders of magnitude, resulting in a 70% heavy sample down to a 0.1% heavy. Because the expected ratio of light to heavy is known, we can compare the measured ratio to assess quantitative accuracy (33). In the DDA dataset, as the sample ratio gets more extreme, there are less proteins detected, due to increasing missingness in the data which makes ratio calculations impossible **(Figure 3A)**. Additionally, as the ratios get more extreme, the measured ratios deviate farther from the expected value. This is especially apparent when comparing the measured ratios for the 70%, 50%, 30%, and 10% ratio samples, where the ratio is reasonably measured, to the 1% ratio sample, which is greatly overestimated by the observed measurements. However, when we measure the same samples by DIA, we have far more measured ratios, due to less missing values **(Figure 3C)**. The number of detected peptides is higher, and while the amount of measured ratios does decrease as the sample ratios get more extreme, there are many more measurements by DIA than by DDA.

**Fig 3.**
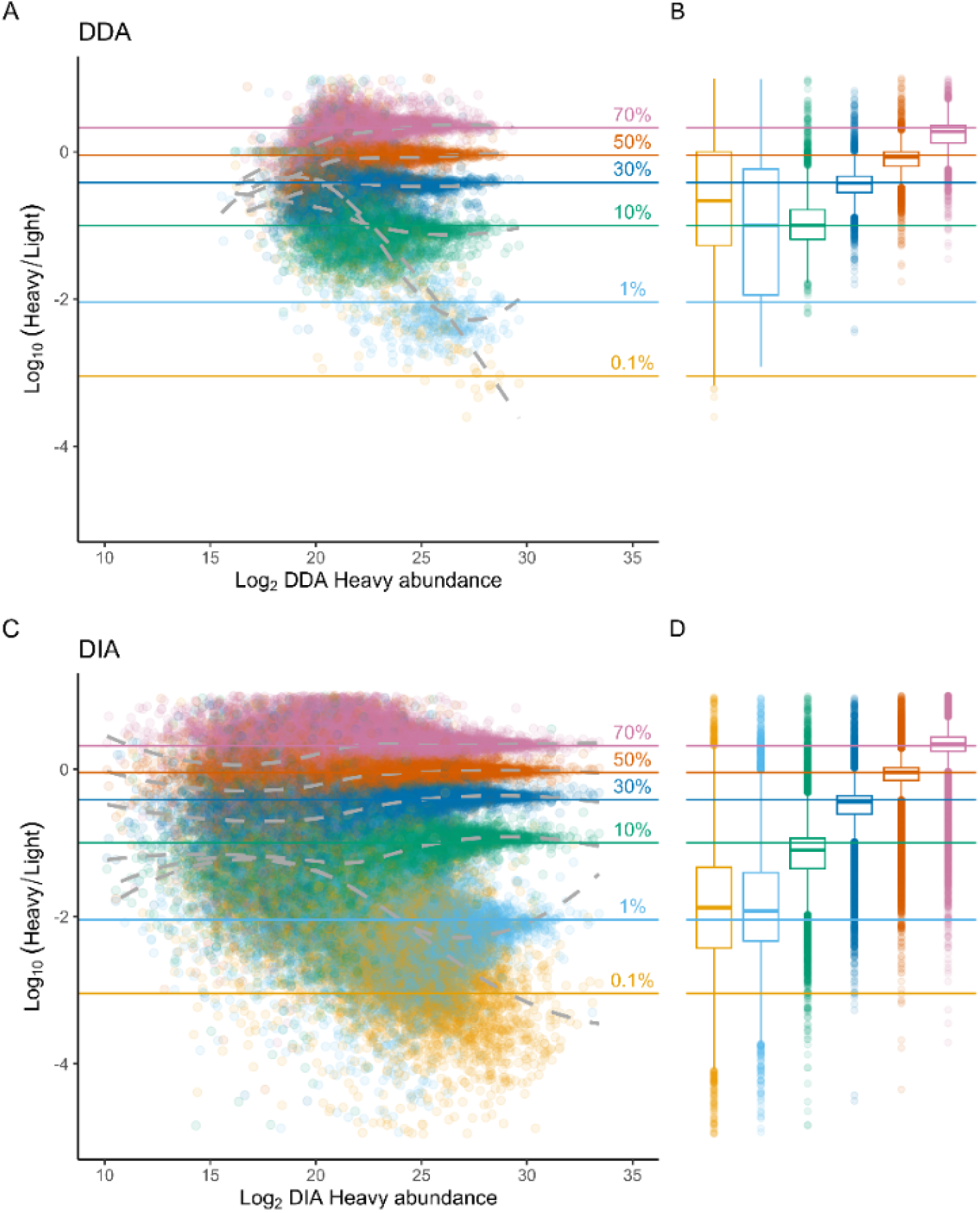
SILAC-DIA improves dynamic range and quantitative accuracy in benchmark experiments. The measured log10(heavy/light) ratios in four dilutions of heavy/light HeLa proteome samples are compared using DDA MS1 quantification with match-between-runs (A, B) enabled versus DIA MS2 quantification (C, D). The samples represent a 70%/30% heavy/light proteome (pink, “0.7”), 50%/50% (orange, “0.5”), 30%/70% (blue, “0.5”), 10/90% (green, “0.1”), 1%/99% (cyan, “0.01”), and a 0.1%/99.9% (yellow, “0.001”). Boxplots depict the first and third quartiles (25th and 75th percentiles) of the Log10(Heavy/Light) values for each sample, with whiskers representing 1.5x the interquartile range.

Further, we see an extended dynamic range by DIA. Although we measure the same highest-abundance peptides, there are many more points extending into the less abundant peptides. Despite lower overall measurements in the most extreme ratio, DIA gained an order of magnitude more sensitivity in peptide ratios than DDA **(Figure 3B, 3D)**. Therefore, it does not appear that isotopic envelope splitting is negatively affecting the SILAC quantifications, at least not as negatively as DDA baseline measurements.

We also measured a similar sample set using *E. coli* on a TripleTOF 6600 system, in which a range of light and heavy *E. coli* ratios (20:1 through 1:20) were assessed for quantitative accuracy **(Supplemental Figure 2)**. Over this 400x range, the ratios measured by SILAC-DIA closely reproduced the expected ratios, demonstrating that SILAC also works for QTOF systems as well as the Orbitrap systems used elsewhere in this work.

To explore whether these qualitative and quantitative benefits translate to biological experiments, we performed a pulse SILAC experiment to determine changes in protein turnover associated with inhibiting the proteasome by bortezomib treatment. Bortezomib is the first FDA-approved proteasome inhibitor for treating cancer (multiple myeloma), and therefore we expect that proteins that are degraded by the ubiquitin-proteasome pathway will have decreased turnover rate (increased half life) in this condition. We dosed cells with either DMSO or bortezomib simultaneously at the time of the SILAC heavy media switch. Using the fractional abundance from each of the ten time points, we fit a nonlinear model to each of the proteins **(Figure 4A, 4B)**. We observed that the distribution of protein turnover models based on the two underlying datasets are overall similar, but that models of longer half lives (decreased turnover) are more disparate compared between DDA and DIA **(Supplemental Figure 3)**. This is in part because, with DIA, ratios roughly an order of magnitude more sensitive can be measured by DIA than by DDA. Additionally, with the DDA data, smaller fractions were more prone to missing values, which is again reflected in these turnover models because there is not enough data at the lower fractions to fit the model. Finally, we note the difference in quantitative dynamic range between the DDA and DIA model distributions, which is also observed in the quantitative benchmarking, recurring again in this application as increased dynamic range in turnover models.

**Figure 4.**
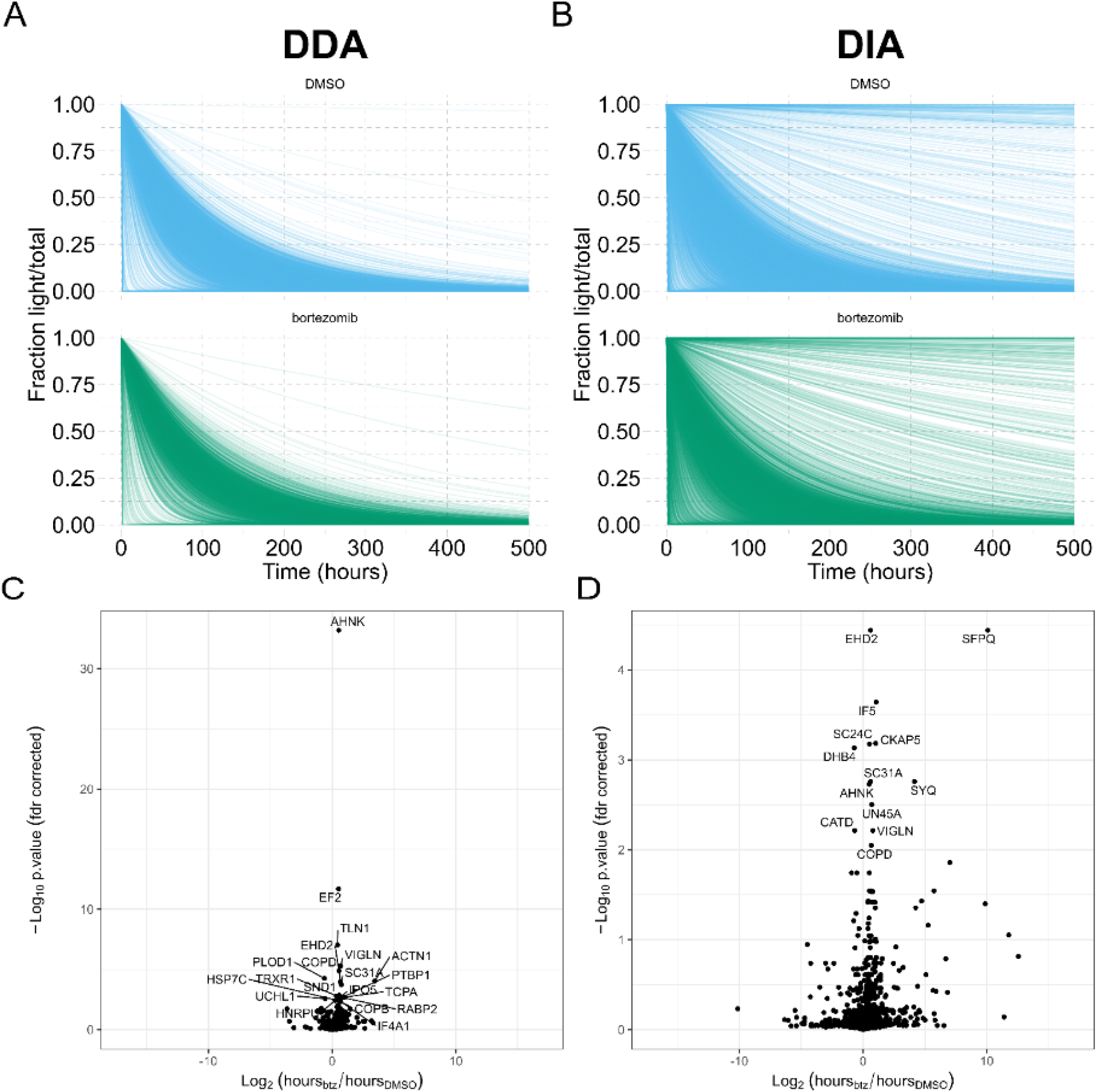
Protein turnover for bortezomib treated cell cultures. **(A, B)** Time is shown horizontally with the fraction of light/total protein vertically. DMSO treatment is shown in blue and bortezomib treatment in green. The shape of these models indicates degradation rate, where proteins with sharply decreasing light fractions are being rapidly degraded, while proteins with more gradual decreases in light fraction are more slowly degraded. **(C, D)** Volcano plots of significantly differential protein half-lives are shown for DDA and DIA analyses of the same samples for DDA (left) and DIA (right).

We compared the differentially degraded proteins as determined using the DDA quantification or the DIA quantification **(Figure 5C, 5D) (Supplemental Data 1)**. A limitation in the workflow approach presented here for pulseSILAC-DIA data processing is that it does not consider any heavy peptides that are not originally present in the light channel, so if there’s some new protein that is present in heavy but not in light, then that protein would not be measured by this method (false negative). Additionally, we have found that a challenge common to all turnover proteomics is the difficulty in calculating protein half-lives and performing differential testing. Here, we perform stringent filtering at the peptide level, only using peptides measured in at least eight of the ten time points, to fit the half-life model. Further, we perform differential testing for proteins with at least three peptides, in order to calculate a mean and variance for each protein half-life, which has excluded many proteins especially from the DIA data analysis. In the future, developing more robust and easy-to-use open-source software for calculating half lives and performing differential testing for turnover experiments would greatly advance the proof-of-concepts shown here.

Although the DDA data tested less proteins for differential degradation, it produced more statistically significant proteins. Specifically, of the 622 proteins tested for differential degradation in the DDA data, 52 were determined statistically significant (q-value < 0.05); of the 1373 proteins tested in the DIA data, 34 were determined statistically significant. The DDA-based significant proteins were estimated from an average of 20 peptides per protein; the DIA-based significant proteins were estimated from an average of 13 peptides per protein. Of the 52 significant proteins by DDA, 3 proteins were detected in the DIA dataset but did not have sufficient peptides to estimate a variance (<3 peptides detected per protein); likewise, of the 34 significant proteins by DIA, 5 were detected by DDA but with only two peptides. It follows that a more robust differential test that isn’t dependent on a minimum of three peptides to estimate variance could improve the agreement between these two methods. At the peptide level, there is a correlation between the half life calculated from the DDA and DIA data **(Supplemental Figure 4)**, suggesting that disagreement between the two datasets arises from rolling peptide values up to protein level. The larger number of significantly differential proteins by DDA is possibly due to the lower number of total tested proteins. Because over twice as many proteins were statistically tested with the DIA dataset, multiple hypothesis correction impacts that dataset more than the DDA dataset.

The two datasets both revealed 12 proteins **(Table 1)** with similar, significant changes to protein half life. By inhibiting the proteasome with bortezomib, we expect that more proteins should be present for a longer time, because they are not degraded by the proteasome. We observe that the majority of these proteins have positive log2 FC, indicating that the protein is present for a longer time (is not degraded) under bortezomib treatment as expected. Of note, the ubiquitin-proteasome pathway is known to degrade HNRNPK (34) and inhibition of the proteasome shows increased levels of HNRNPK, supporting our results. Additionally, the 20S proteasome has been shown to degrade EIF3A and IF4A1/EIF4A-1 (35), which we also observe here in the form of decreased degradation under proteasome inhibition. We also note that three proteins appear to be degraded faster (or synthesized more) under bortezomib treatment. Two of these three proteins, DHB4 and CATD, are also known to have cleaved forms, which may not be detected by the approach performed here as it only considers the protein level. Increased half life of these proteins may indicate that, after time zero and treatment with bortezomib, the protein can no longer be cleaved into its products. More elegant models, either for protein quantification or for differential turnover at the peptide level, would potentially be useful for such systems as bortezomib treatment, which may affect not only global protein abundances but also cleavage isoforms. The decrease in Cathepsin D (CATD) degradation half-life measured here could also be explained by an increased synthesis rate, as cathepsin D is an aspartyl protease and could be upregulated to counter the cell’s inability to degrade proteins via the bortezomib-inhibited ubiquitin-proteasome pathway. Increases in CATD due to either increased transcription rates or alternative processing of the pro-CATD gene product have been linked to breast cancer metastasis (36).

**Table 1.**
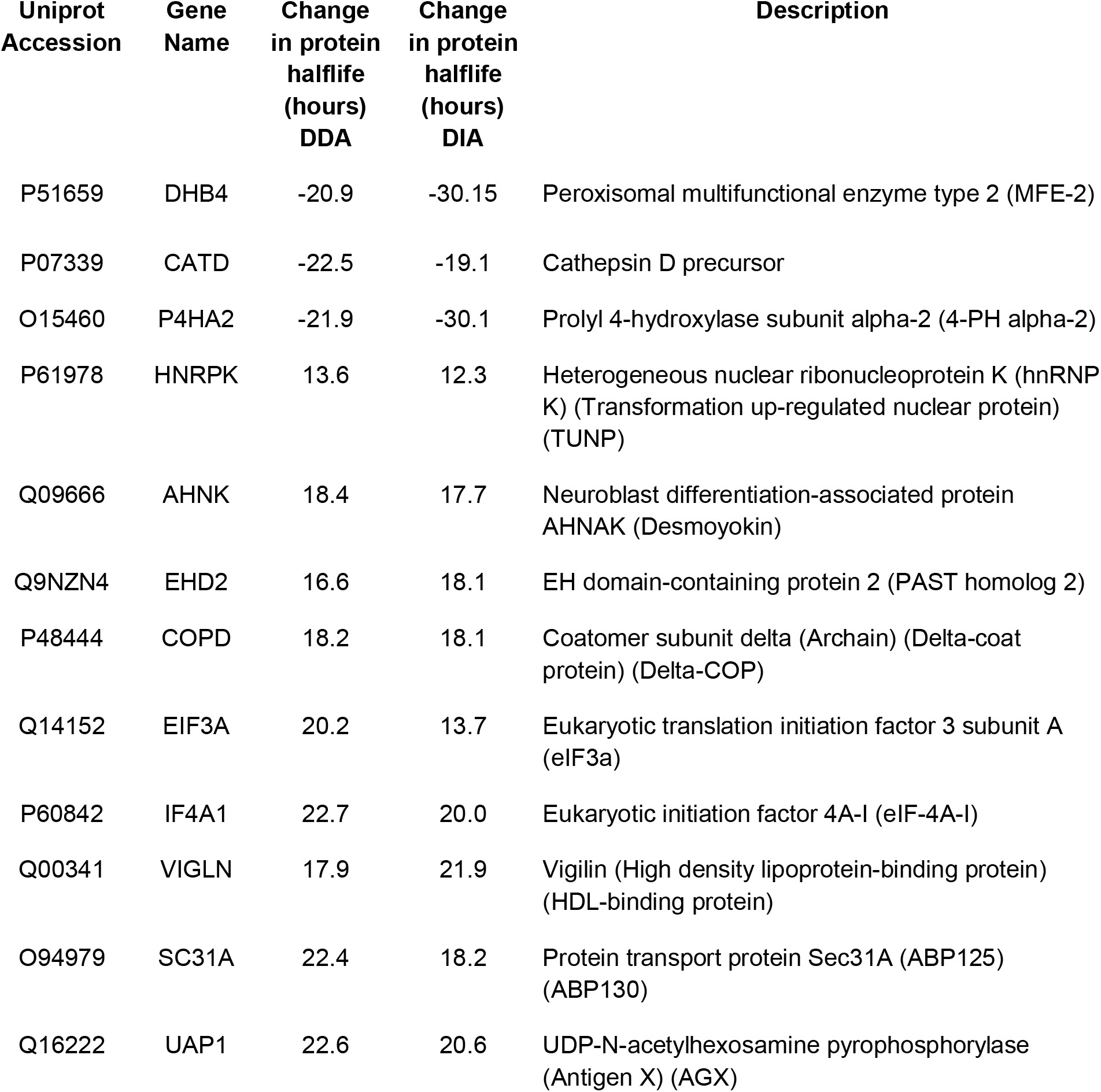
Significantly differentially degraded proteins (bortezomib vs DMSO) found by both DDA and DIA half life estimation. The twelve shared proteins found to be differential using both the DDA and the DIA datasets are shown along with the log2 fold change (bortezomib/DMSO) calculated by each dataset.

## CONCLUSIONS

The experiments described here validate the combination of SILAC together with DIA, demonstrating that the theoretical concern for isotopic envelope splitting detrimentally affecting peptide detection and quantification is not observed in empirical data. Using benchmarking data sets, we show that conventional DIA methods can be used for SILAC samples and that narrower DIA isolation window methods outperform wider window schemes in peptide detection; but multiplex SILAC-DIA does not detect as many peptides as multiplex SILAC-DDA. Less peptides are detected in the 100% light and 100% heavy samples when the search space contains both light and heavy precursors likely due to the search algorithm (Walnut) having difficulty with handling the doubled search space. However, that limitation is overcome in the 50/50% sample because then the DDA approach is limited in that half the topX DDA precursor peaks are redundant isotope pairs.

SILAC-DIA truly outperforms not in peptide detections but rather in peptide quantification, achieving an order of magnitude more depth in quantitative accuracy compared to conventional SILAC-DDA. We also demonstrate that SILAC-DIA quantification matches expected ratios not only for Orbitrap systems using fixed and staggered windows, but for QTOF instruments using variable width windowing as well, showing that our observations are generalizable across instrumentation and DIA isolation schemes. Therefore, while SILAC-DIA could potentially be used for multiplex experiments, where a light and heavy sample are combined at an equimolar ratio, it is our recommendation that the best application for SILAC-DIA is pSILAC turnover experiments, where more precise and accurate quantification at extreme ratios can be used to fit more sensitive turnover models.

As our approach for processing pSILAC-DIA described here is readily performed with freely-available open-source software and is detailed with a full tutorial (**Supplementary Note 1**), and with the sustained interest in using DIA approaches, we anticipate this workflow becoming a popular choice for pulse SILAC-based protein turnover experiments.

## Supporting information

Supplemental Data 1. Statistical testing results for differential protein half-lives under bortezomib treatment using DDA and DIA quantifications.

Supplemental Note 1. Tutorial for analyzing pulseSILAC-DIA data with Prosit, EncyclopeDIA and Skyline.

Supplemental Table 3. Data-independent acquisition methods parameters and windowing schemes used in this work.

## ACKNOWLEDGEMENTS

This work was supported by the National Institutes of Health R01GM110174 and P01AG031862 and by Celgene to BAG; and by the National Institutes of Health/National Cancer Institute T32CA009140 to LKP. B.S. was supported by a Nathan Shock Center Pilot Award (University of Washington). Computational resources were supported by NIH Project Grant S10OD023592. We additionally acknowledge the support of instrumentation from the NCRR shared instrumentation grant 1S10 OD016281 to the Buck Institute.

## SUPPORTING INFORMATION

**Supplemental Table 1.** Serial dilution scheme used for constructing pulseSILAC calibration curves.

**Supplemental Table 2.** Serial dilution scheme used for constructing E.coli heavy/light SILAC ratio samples.

**Supplemental Table 3.** Data-independent acquisition methods parameters and windowing schemes used in this work.

**Supplemental Figure 1.** Simulation of precursor isotope envelopes of SILAC light and heavy peptides.

**Supplemental Figure 2.** SILAC-DIA quantification closely reproduces expected ratios of light/heavy *E. coli* mixtures.

**Supplemental Figure 3.** Distribution of protein half lives under DMSO and bortezomib, as modeled using DDA and DIA data.

**Supplemental Figure 4.** Correlation of peptide half lives as calculated by DDA and DIA.

**Supplemental Figure 5.** Distribution of significant protein half lives as assessed by DDA and DIA.

**Supplemental Data 1.** Statistical testing results for differential protein half-lives under bortezomib treatment using DDA and DIA quantifications.

**Supplemental Note 1.** Tutorial for analyzing pulseSILAC-DIA data with Prosit+EncyclopeDIA and Skyline.

