## Supplemental Figures and Tables for "Improved SILAC quantification with data independent acquisition to investigate bortezomib-induced protein degradation"

**Supplemental Figure 1.** Simulation of precursor isotope envelopes of SILAC light and heavy peptides.

**Supplemental Figure 2.** SILAC-DIA quantification closely reproduces expected ratios of light/heavy *E. coli* mixtures.

**Supplemental Figure 3.** Distribution of protein half lives under DMSO and bortezomib, as modeled using DDA and DIA data.

**Supplemental Figure 4.** Correlation of peptide half lives as calculated by DDA and DIA.

**Supplemental Figure 5.** Distribution of significant protein half lives as assessed by DDA and DIA.

**Supplemental Table 1.** Serial dilution scheme used for constructing pulseSILAC calibration curves.

**Supplemental Table 2.** Serial dilution scheme used for constructing E.coli heavy/light SILAC ratio samples.

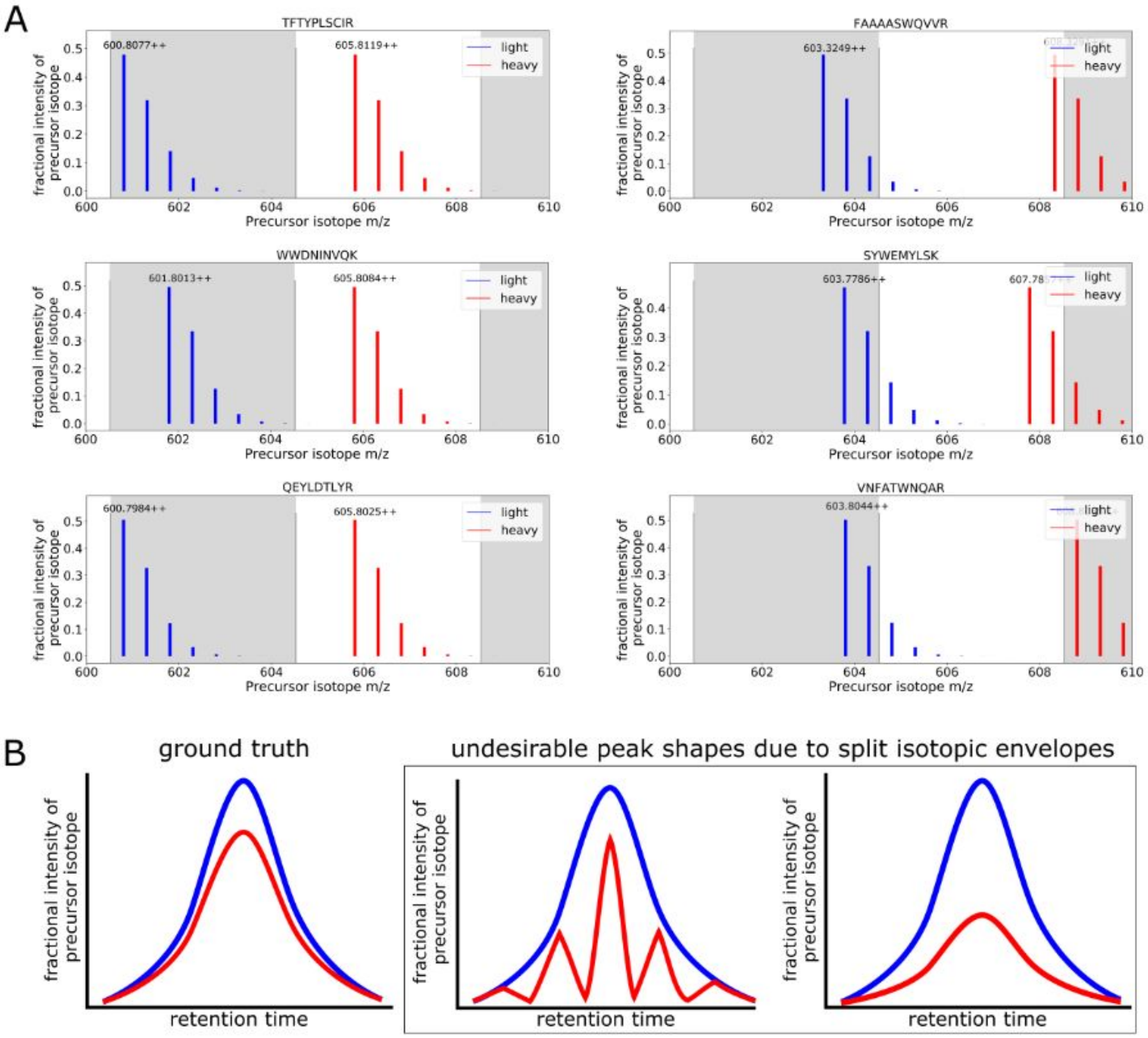


**Supplemental Figure 1. Simulation of precursor isotope envelopes of SILAC light and heavy peptides. (A)** Six peptides generated from an in silico digestion of the human reference proteome are shown with their light precursor isotopic envelope (blue) and their respective heavy arginine/lysine SILAC isotopic envelope (red). Proposed DIA isolation window boundaries (gray) bound each precursor within a single window (left column), but split the isotopic envelopes of other precursor pairs (right column). **(B)** A ground truth chromatogram for a hypothetical light and heavy precursor pair is shown (left) along with two undesirable chromatograms (right) which may arise due to the heavy precursor having its isotopic envelope split across DIA isolation windows in situations with staggered windows or fixed-width windows.

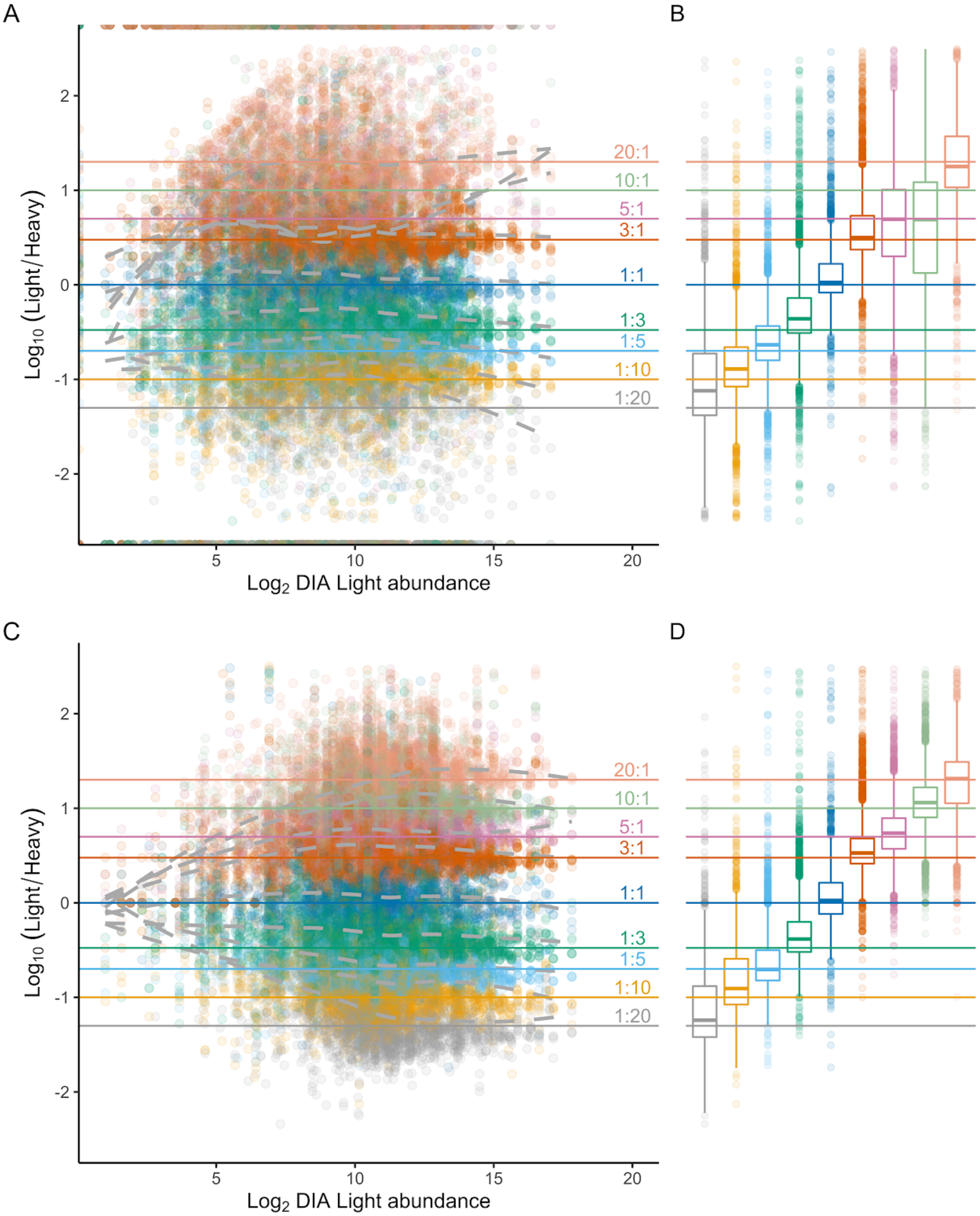


**Supplemental Figure 2. SILAC-DIA quantification closely reproduces expected ratios of light/heavy *E.coli* mixtures.** The measured log10(light/heavy) ratios in nine dilutions of heavy/light *E.coli* proteome samples are compared using SWATH MS2 quantification. The samples represent dilutions of light *E.coli* into heavy *E.coli* with ratios between 20:1 through 1:20, a 400x range.

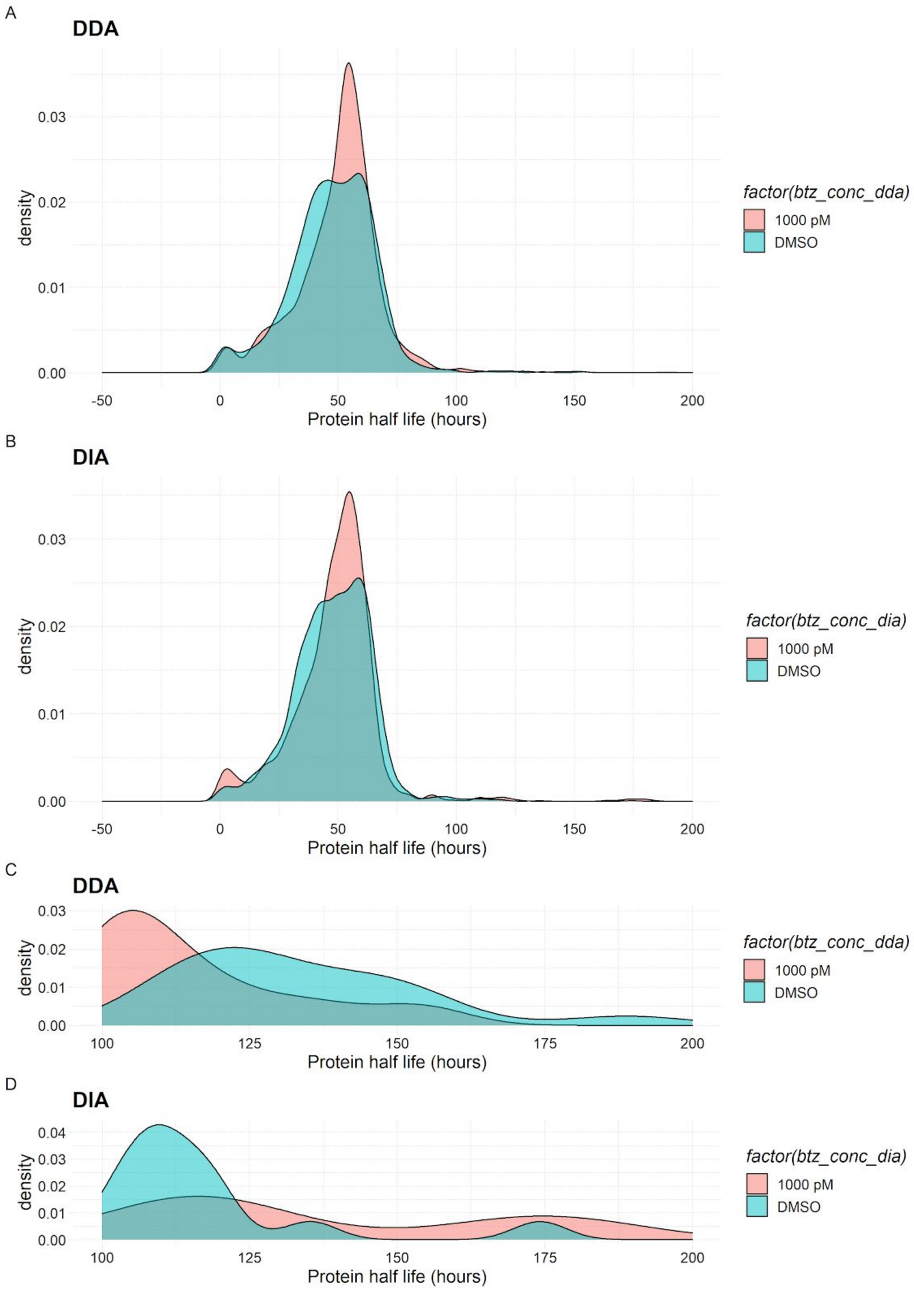


**Supplemental Figure 3. Distribution of protein half lives under DMSO and bortezomib, as modeled using DDA and DIA data.** Protein half lives based on >2 peptides for each protein are shown as a density distribution for bortezomib treatment (1000 pM) and DMSO control, with the half lives as calculated by DDA and DIA. **(A,B)** The overall distribution of half life values calculated by DDA and DIA are quite similar. **(C,D)** The more extreme half life values (100-200 hours) are shown to visualize the differences in model sensitivity.

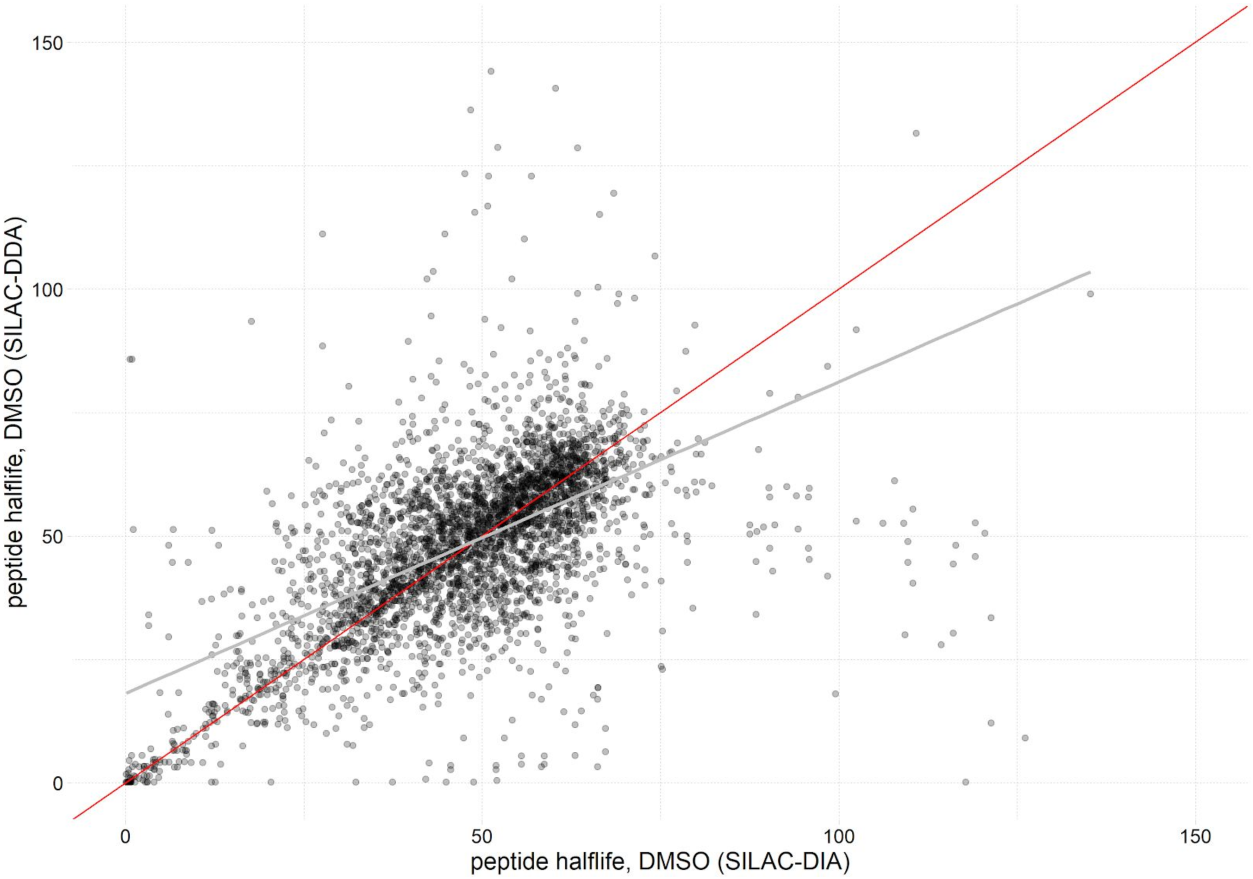


**Supplemental Figure 4. Correlation of peptide half lives as calculated by DDA and DIA.** Each peptide half life is shown with the value as calculated by DIA against the value as calculated by DDA data. A line of equality (red) is plotted along with a line of best fit (gray). The <1 slope shows that a half life estimated by DDA tends to be smaller (shorter) than a half life estimated by DIA.

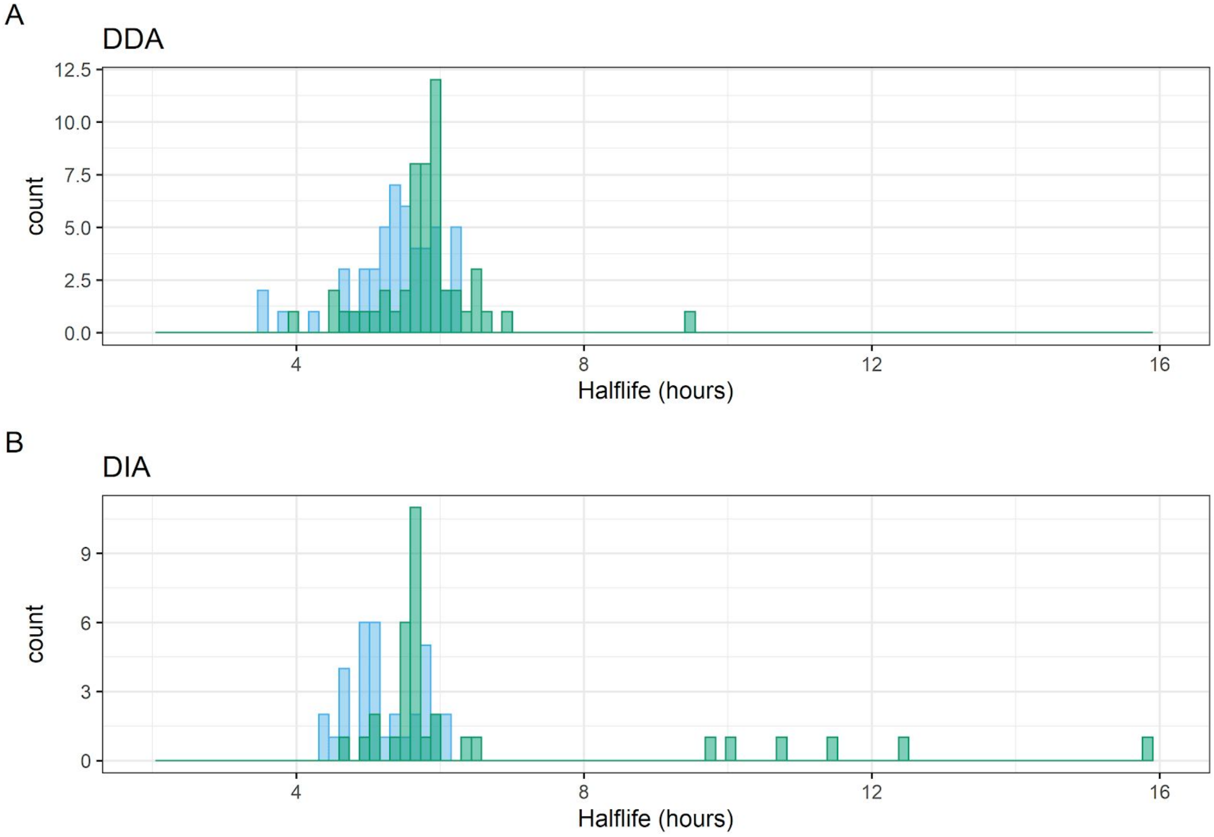


**Supplemental Figure 5. Distribution of significant protein half lives as assessed by DDA and DIA.** The differentially degraded proteins as determined by DDA-based models **(A)** and by DIA-based models **(B)** are shown for bortezomib treatment (green) and DMSO control (blue). Overall, there are more significantly degraded proteins as reported by the DDA-based models, but the DIA-based models include more extreme half life values over 8 hours.

| Point | 100% Heavy sample (ul) | 100% Light sample (ul) | Sample to use for serial dilution | Serial dilution (ul) | Light (ul) | frx dilution Heavy |
|---|---|---|---|---|---|---|
| A | 100 | 0 | | | | 1 |
| B | 70 | 30 | | | | 0.7 |
| C | 50 | 50 | | | | 0.5 |
| D | 30 | 70 | | | | 0.3 |
| E | 10 | 90 | | | | 0.1 |
| F | | | B | 10 | 90 | 0.07 |
| G | | | C | 10 | 90 | 0.05 |
| H | | | D | 10 | 90 | 0.03 |
| I | | | E | 10 | 90 | 0.01 |
| J | | | F | 10 | 90 | 0.007 |
| K | | | G | 10 | 90 | 0.005 |
| L | | | H | 10 | 90 | 0.003 |
| M | | | I | 10 | 90 | 0.001 |
| N | 0 | 100 | | | | 0 |

**Supplemental Table 1. Dilution scheme for preparing mixture samples of HeLa SILAC light and heavy.**

| Sample Name | Light E.coli | Heavy E.coli | Percent light | Percent heavy |
|---|---|---|---|---|
| 20:1 | 20 | 1 | 0.95 | 0.05 |
| 10:1 | 10 | 1 | 0.91 | 0.09 |
| 5:1 | 5 | 1 | 0.83 | 0.17 |
| 3:1 | 3 | 1 | 0.75 | 0.25 |
| 1:1 | 1 | 1 | 0.50 | 0.50 |
| 1:3 | 1 | 3 | 0.25 | 0.75 |
| 1:5 | 1 | 5 | 0.17 | 0.83 |
| 1:10 | 1 | 10 | 0.09 | 0.91 |
| 1:20 | 1 | 20 | 0.05 | 0.95 |

**Supplemental Table 2. Dilution scheme for preparing mixture samples of E.coli SILAC light and heavy.**
