## Supplemental Note 1. Tutorial for analyzing pulseSILAC-DIA data with Prosit, EncyclopeDIA and Skyline. for "Improved SILAC quantification with data independent acquisition to investigate bortezomib-induced protein degradation"

### Supplementary Note 1: Analyzing pulseSILAC-DIA data with Prosit+EncyclopeDIA and Skyline

This tutorial is a practical guide for how to use the Encyclopedia software suite to build. . In this tutorial, we will detail how we analyzed the pulse SILAC-DIA data using EncyclopeDIA to first build a library of detected endogenous peptides, then using Skyline to pair the endogenous precursors to their "heavy" SILAC counterpart and extracting their chromatograms for quantification.

**SUMMARY: Four steps for pulseSILAC-DIA analysis**

1. Convert .raw files to .mzML using MSConvert
    **Not covered here, please see Pino et al 2020 Supplementary Note 1 (https://doi.org/10.1074/mcp.P119.001913) for a detailed tutorial.*
2. Build time point zero library using Prosit and EncyclopeDIA
3. Search pulsed data with library from step 2 using EncyclopeDIA
4. Extract light/heavy chromatograms using Skyline

***Citations***

MSconvert (https://www.nature.com/articles/nbt.2377)
    A cross-platform toolkit for mass spectrometry and proteomics. Chambers MC et al. *Nat Biotech* 30, 918-920 (2012). doi.org/10.1038/nbt.2377

Encyclopedia (https://www.nature.com/articles/s41467-018-07454-w)
    Chromatogram libraries improve peptide detection and quantification by data independent acquisition mass spectrometry. Searle BC et al. *Nat Comm* 9, 5128 (2018). doi.org/10.1038/s41467-018-07454-w

Prosit+Encyclopedia (https://www.nature.com/articles/s41467-020-15346-1)
    Generating high quality libraries for DIA MS with empirically corrected peptide predictions. Searle BC et al. *Nat Commun* 11, 1548 (2020). https://doi.org/10.1038/s41467-020-15346-1

You will need:

- MSConvert from Proteowizard: *Windows only!*
    - http://proteowizard.sourceforge.net/download.html
- EncyclopeDIA suite (*.jar file): *command line and cross-platform GUI*
    - https://bitbucket.org/searleb/encyclopedia/wiki/Home
- Skyline: *Windows only!*
    - https://skyline.ms/project/home/software/Skyline/begin.view

To exactly replicate the results here, you will also need:

- RAW DIA data files from the tutorial bortezomib dataset (PXD022659)
- Ready-made predicted spectral library in *.dlib format and accompanying FASTA of the Uniprot human reference proteome (reviewed; 20,350 entries) available on the Prosit Libraries website

### BUILD THE POOL OF PEPTIDES PRESENT AT TIME ZERO

1. Launch EncyclopeDIA. Under “Parameters:” click the “Library:” Edit button. Navigate to and select your Prosit predicted library (in DLIB format). Then, click the “Background:” Edit button to navigate to and select the corresponding FASTA for that Prosit library.

   ***! NOTE**: for a detailed tutorial on generating custom Prosit libraries, please see the Supplementary Info for Searle et al 2020*

   ***! NOTE**: Prosit libraries for common organisms (human and yeast) can also be found on the Prosit website here.*

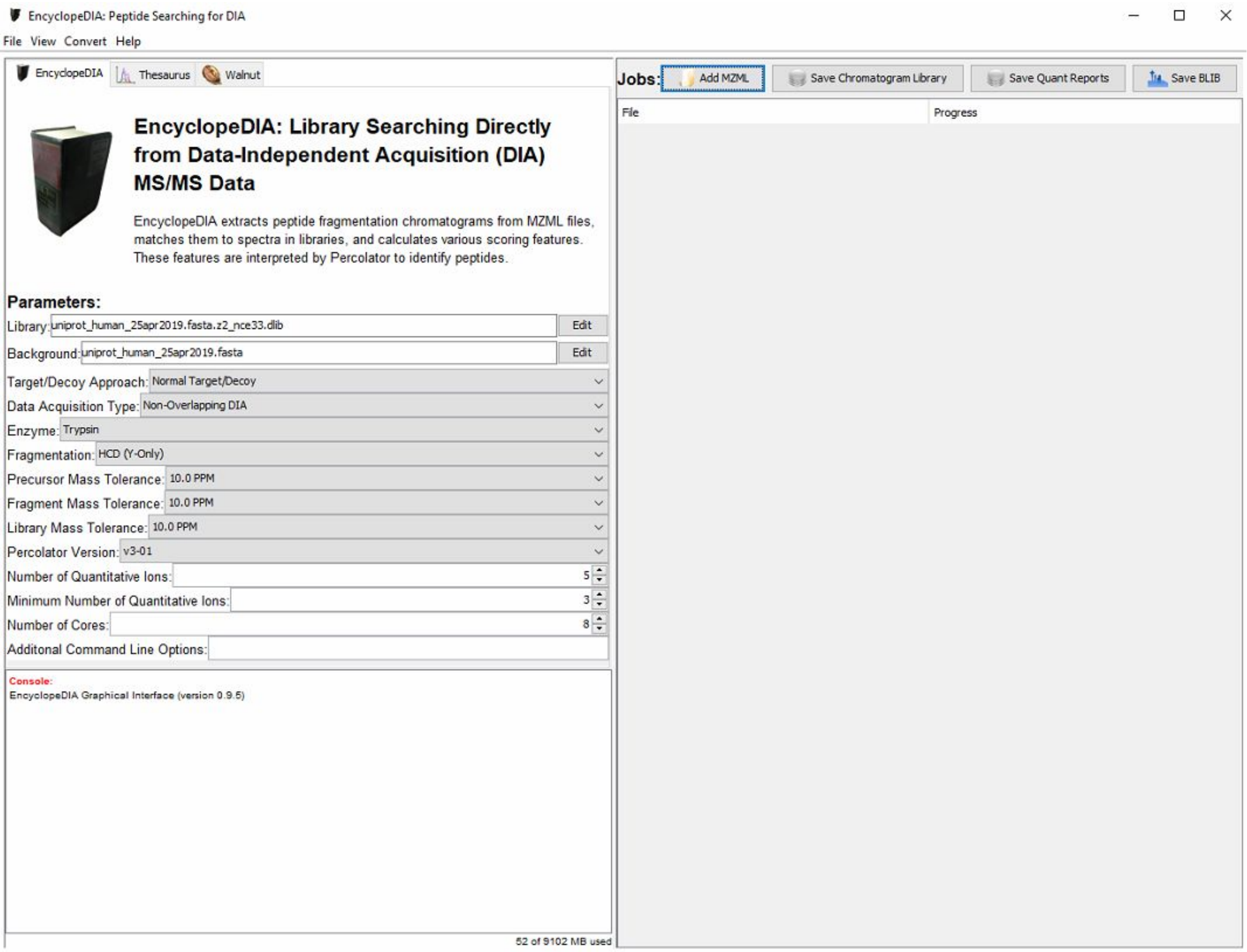


2. At the top next to “Jobs:”, click “Add MZML” to select the narrow window, gas phase fractionated (GPF) MZML files for your experiment.

***! NOTE**: The GPF library should be composed of the "time zero" sample from your pulse SILAC experiment. In other words, this pooled library sample should be composed entirely of endogenous/light peptides prior to incorporation of heavy SILAC amino acids.*

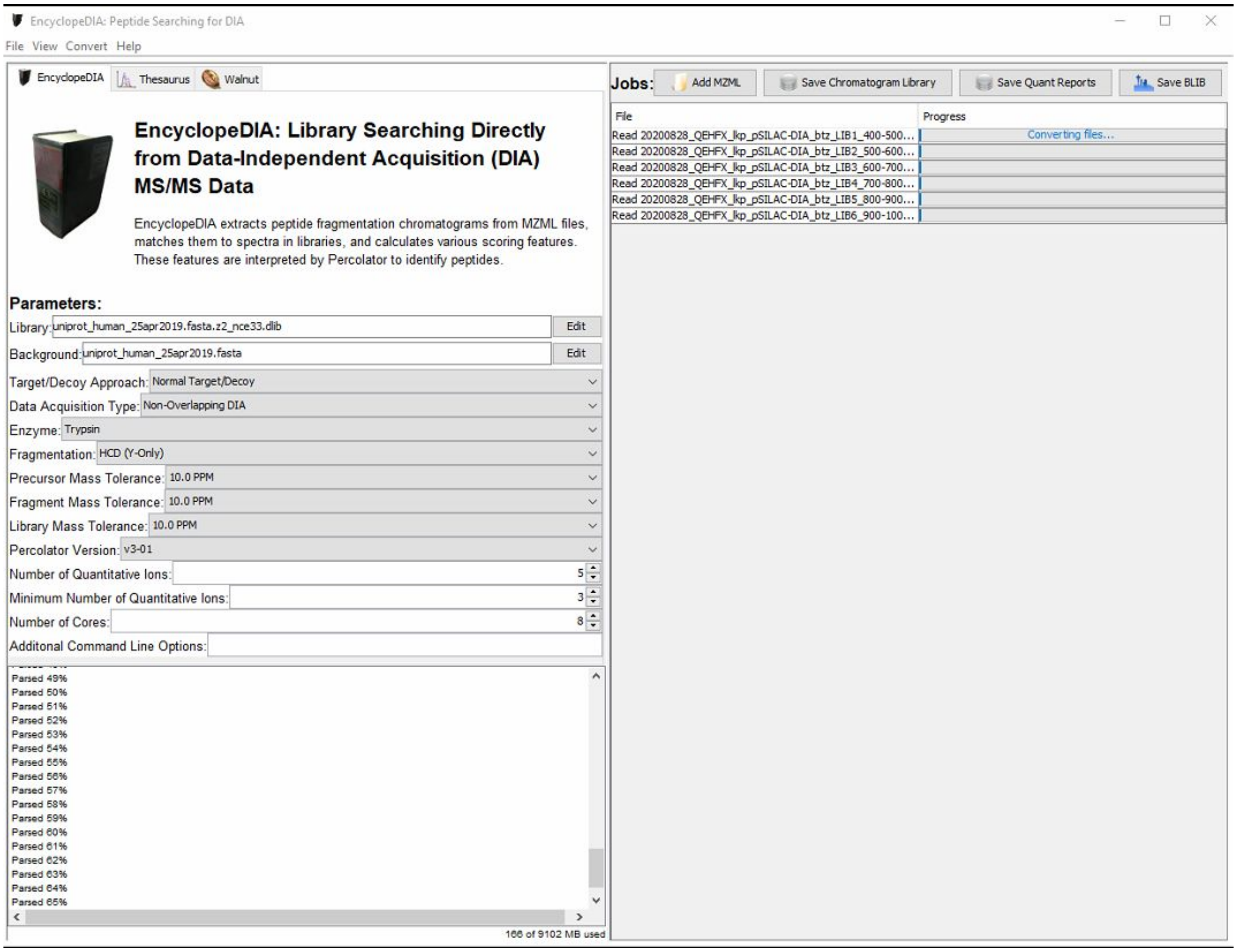


3. After all GPF library files have completed, click "Save Chromatogram Library" to perform the final FDR filtering and save the ELIB.

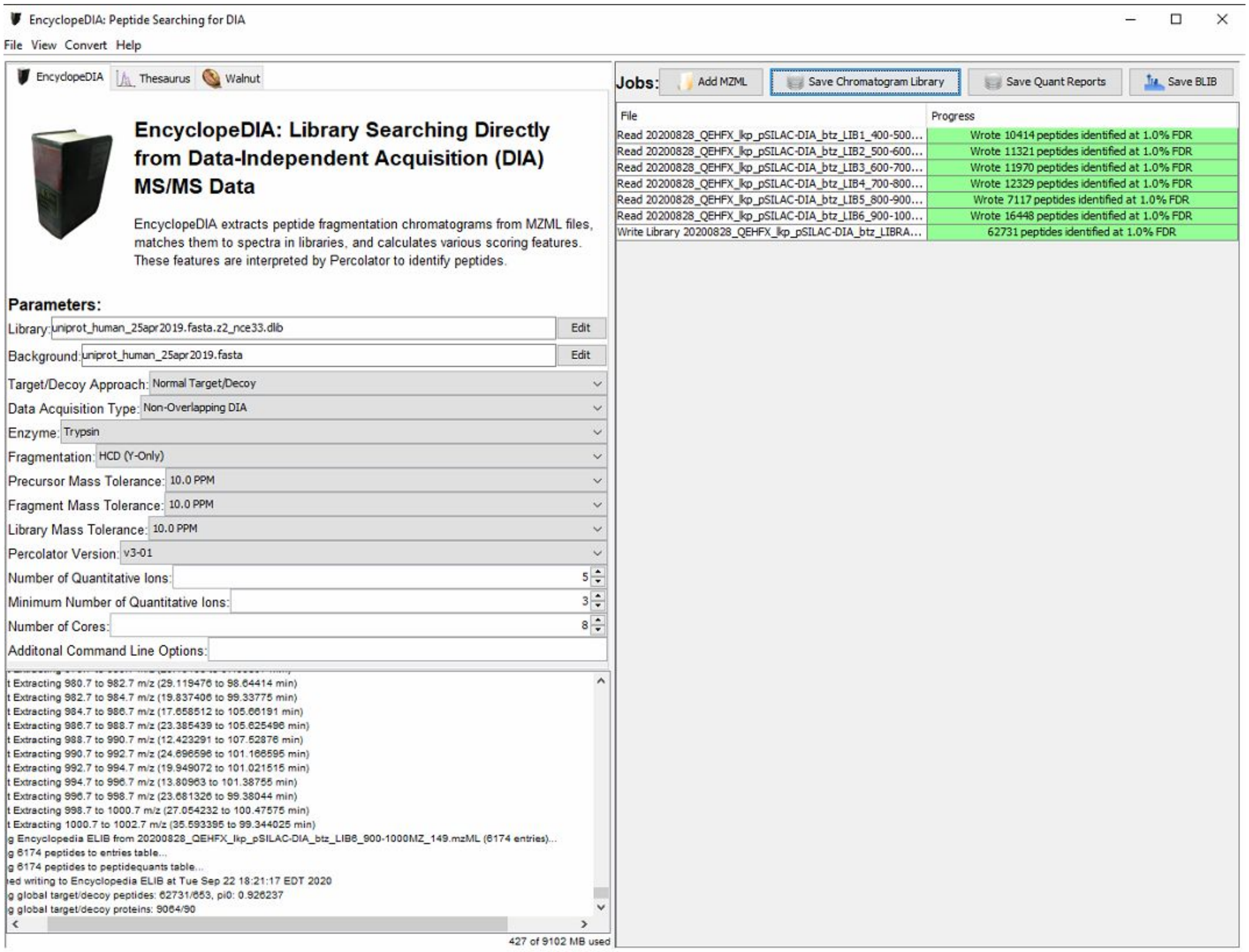
EncyclopeDIA: Peptide Searching for DIA
File View Convert Help
EncyclopeDIA
Thesaurus
Walnut
EncyclopeDIA: Library Searching Directly from Data-Independent Acquisition (DIA) MS/MS Data
EncyclopeDIA extracts peptide fragmentation chromatograms from MZML files, matches them to spectra in libraries, and calculates various scoring features. These features are interpreted by Percolator to identify peptides.
Parameters:
Library: uniprot_human_25apr2019.fasta.z2_nce33.dlib
Edit
Background: uniprot_human_25apr2019.fasta
Edit
Target/Decoy Approach: Normal Target/Decoy
Data Acquisition Type: Non-Overlapping DIA
Enzyme: Trypsin
Fragmentation: HCD (Y-Only)
Precursor Mass Tolerance: 10.0 PPM
Fragment Mass Tolerance: 10.0 PPM
Library Mass Tolerance: 10.0 PPM
Percolator Version: v3-01
Number of Quantitative Ions: 5
Minimum Number of Quantitative Ions: 3
Number of Cores: 8
Additonal Command Line Options:
427 of 9102 MB used
Jobs:
Add MZML
Save Chromatogram Library
Save Quant Reports
Save BLIB
File
Progress
Read 20200828_QEHFX_lkp_pSILAC-DIA_btz_LIB1_400-500...
Wrote 10414 peptides identified at 1.0% FDR
Read 20200828_QEHFX_lkp_pSILAC-DIA_btz_LIB2_500-600...
Wrote 11321 peptides identified at 1.0% FDR
Read 20200828_QEHFX_lkp_pSILAC-DIA_btz_LIB3_600-700...
Wrote 11970 peptides identified at 1.0% FDR
Read 20200828_QEHFX_lkp_pSILAC-DIA_btz_LIB4_700-800...
Wrote 12329 peptides identified at 1.0% FDR
Read 20200828_QEHFX_lkp_pSILAC-DIA_btz_LIB5_800-900...
Wrote 7117 peptides identified at 1.0% FDR
Read 20200828_QEHFX_lkp_pSILAC-DIA_btz_LIB6_900-100...
Wrote 16448 peptides identified at 1.0% FDR
Write Library 20200828_QEHFX_lkp_pSILAC-DIA_btz_LIBRA...
62731 peptides identified at 1.0% FDR

#### SEARCH PULSED DATA WITH LIBRARY FROM TIME ZERO

4. Close and relaunch EncyclopeDIA to clear the cache. Under “Parameters:” click the “Library:” Edit button. Navigate to and select the chromatogram library you created in Step 3 above (in ELIB format). Then, click the “Background:” Edit button to navigate to and select the corresponding FASTA for the original Prosit library.

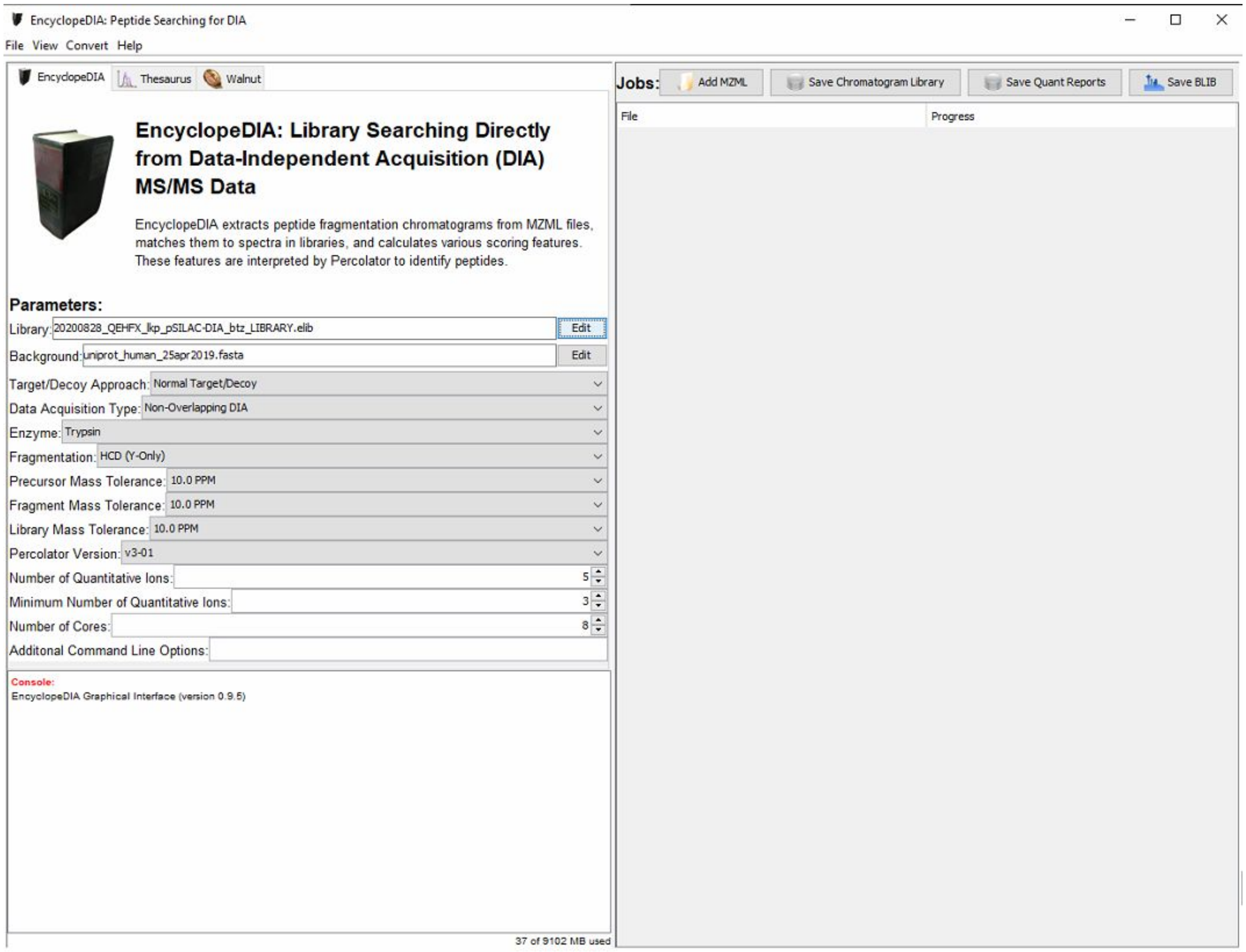


5. At the top next to “Jobs:”, click “Add MZML” to select the wide window, single-shot MZML files for your experiment.

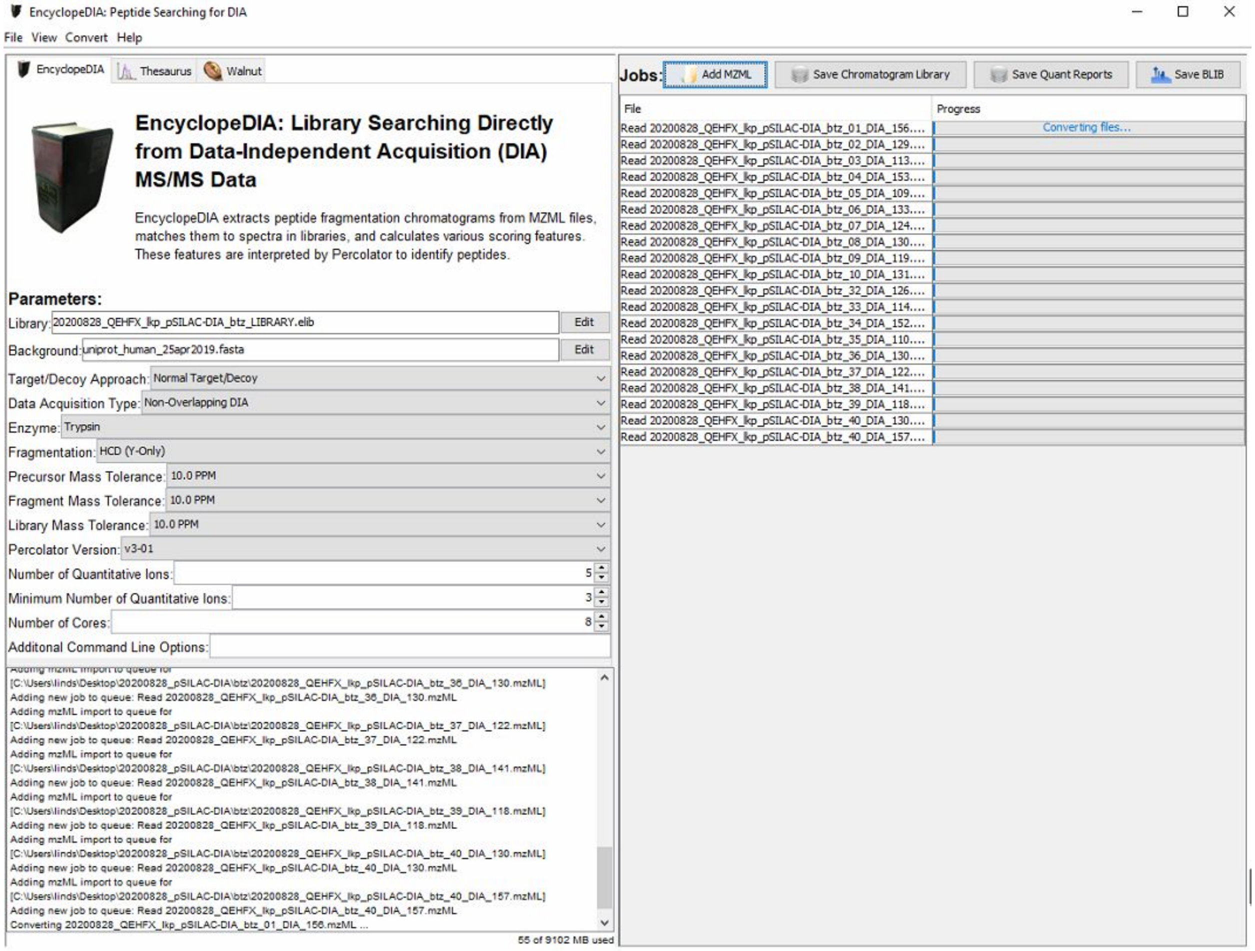


6. After all wide window, single-shot files have completed, click “Save Quant Report” to perform the global retention time alignment, fragment ion refinement, and FDR filtering; and save the ELIB.

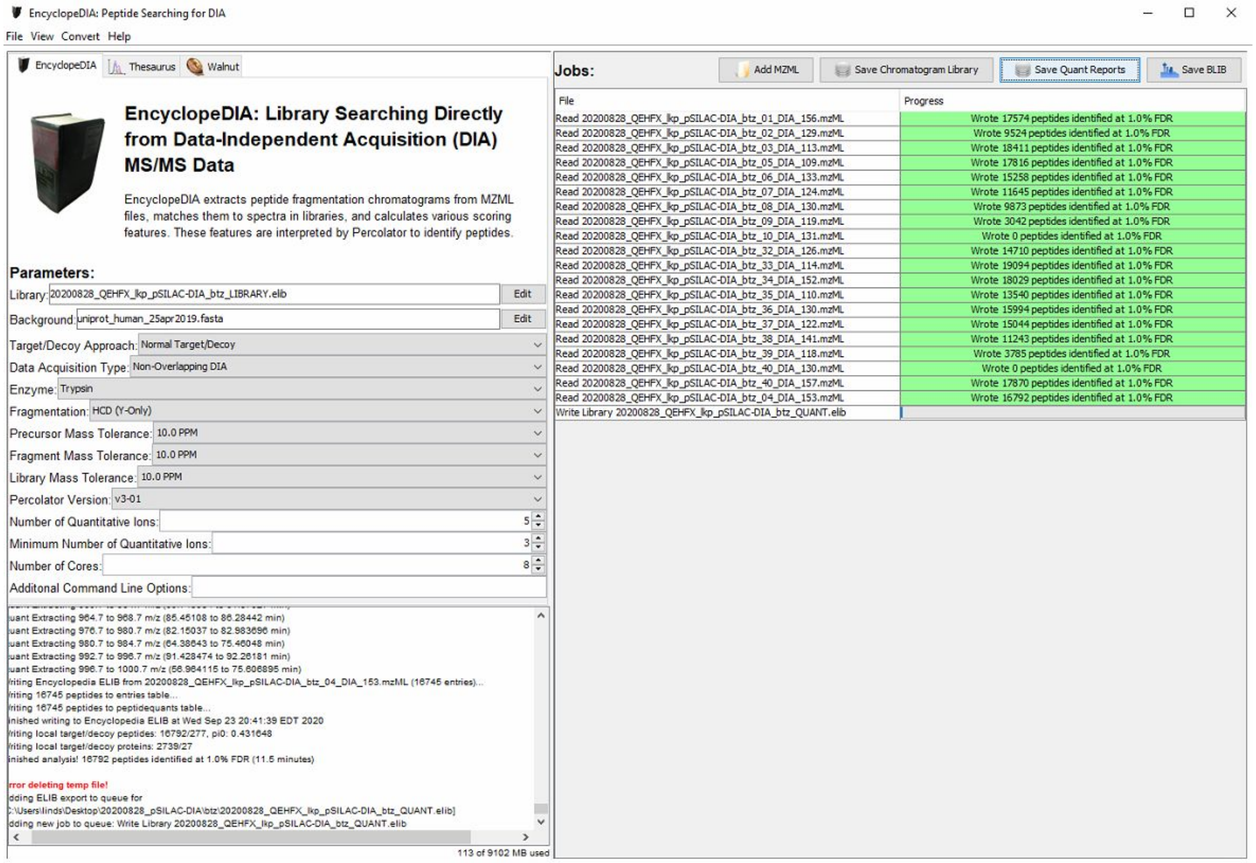
EncyclopeDIA: Peptide Searching for DIA
File View Convert Help
EncyclopeDIA
Thesaurus
Walnut
EncyclopeDIA: Library Searching Directly from Data-Independent Acquisition (DIA) MS/MS Data
EncyclopeDIA extracts peptide fragmentation chromatograms from MZML files, matches them to spectra in libraries, and calculates various scoring features. These features are interpreted by Percolator to identify peptides.
Parameters:
Library: 20200828_QEHFX_lkp_pSILAC-DIA_btz_LIBRARY.elib
Edit
Background: uniprot_human_25apr2019.fasta
Edit
Target/Decoy Approach: Normal Target/Decoy
Data Acquisition Type: Non-Overlapping DIA
Enzyme: Trypsin
Fragmentation: HCD (Y-Only)
Precursor Mass Tolerance: 10.0 PPM
Fragment Mass Tolerance: 10.0 PPM
Library Mass Tolerance: 10.0 PPM
Percolator Version: v3-01
Number of Quantitative Ions: 5
Minimum Number of Quantitative Ions: 3
Number of Cores: 8
Additonal Command Line Options:
uant Extracting 964.7 to 968.7 m/z (85.45108 to 86.28442 min)
uant Extracting 976.7 to 980.7 m/z (82.15037 to 82.983696 min)
uant Extracting 980.7 to 984.7 m/z (64.38643 to 75.46048 min)
uant Extracting 992.7 to 996.7 m/z (91.428474 to 92.26181 min)
uant Extracting 996.7 to 1000.7 m/z (56.964115 to 75.606895 min)
/riting Encyclopedia ELIB from 20200828_QEHFX_lkp_pSILAC-DIA_btz_04_DIA_153.mzML (16745 entries)...
/riting 16745 peptides to entries table...
/riting 16745 peptides to peptidequants table...
inished writing to Encyclopedia ELIB at Wed Sep 23 20:41:39 EDT 2020
/riting local target/decoy peptides: 16792/277, pi0: 0.431648
/riting local target/decoy proteins: 2739/27
inished analysis! 16792 peptides identified at 1.0% FDR (11.5 minutes)
rror deleting temp file!
dding ELIB export to queue for
:\Users\linds\Desktop\20200828_pSILAC-DIA\btz\20200828_QEHFX_lkp_pSILAC-DIA_btz_QUANT.elib]
dding new job to queue: Write Library 20200828_QEHFX_lkp_pSILAC-DIA_btz_QUANT.elib
113 of 9102 MB used
Jobs:
Add MZML
Save Chromatogram Library
Save Quant Reports
Save BLIB
File
Progress
Read 20200828_QEHFX_lkp_pSILAC-DIA_btz_01_DIA_156.mzML
Wrote 17574 peptides identified at 1.0% FDR
Read 20200828_QEHFX_lkp_pSILAC-DIA_btz_02_DIA_129.mzML
Wrote 9524 peptides identified at 1.0% FDR
Read 20200828_QEHFX_lkp_pSILAC-DIA_btz_03_DIA_113.mzML
Wrote 18411 peptides identified at 1.0% FDR
Read 20200828_QEHFX_lkp_pSILAC-DIA_btz_05_DIA_109.mzML
Wrote 17816 peptides identified at 1.0% FDR
Read 20200828_QEHFX_lkp_pSILAC-DIA_btz_06_DIA_133.mzML
Wrote 15258 peptides identified at 1.0% FDR
Read 20200828_QEHFX_lkp_pSILAC-DIA_btz_07_DIA_124.mzML
Wrote 11645 peptides identified at 1.0% FDR
Read 20200828_QEHFX_lkp_pSILAC-DIA_btz_08_DIA_130.mzML
Wrote 9873 peptides identified at 1.0% FDR
Read 20200828_QEHFX_lkp_pSILAC-DIA_btz_09_DIA_119.mzML
Wrote 3042 peptides identified at 1.0% FDR
Read 20200828_QEHFX_lkp_pSILAC-DIA_btz_10_DIA_131.mzML
Wrote 0 peptides identified at 1.0% FDR
Read 20200828_QEHFX_lkp_pSILAC-DIA_btz_32_DIA_126.mzML
Wrote 14710 peptides identified at 1.0% FDR
Read 20200828_QEHFX_lkp_pSILAC-DIA_btz_33_DIA_114.mzML
Wrote 19094 peptides identified at 1.0% FDR
Read 20200828_QEHFX_lkp_pSILAC-DIA_btz_34_DIA_152.mzML
Wrote 18029 peptides identified at 1.0% FDR
Read 20200828_QEHFX_lkp_pSILAC-DIA_btz_35_DIA_110.mzML
Wrote 13540 peptides identified at 1.0% FDR
Read 20200828_QEHFX_lkp_pSILAC-DIA_btz_36_DIA_130.mzML
Wrote 15994 peptides identified at 1.0% FDR
Read 20200828_QEHFX_lkp_pSILAC-DIA_btz_37_DIA_122.mzML
Wrote 15044 peptides identified at 1.0% FDR
Read 20200828_QEHFX_lkp_pSILAC-DIA_btz_38_DIA_141.mzML
Wrote 11243 peptides identified at 1.0% FDR
Read 20200828_QEHFX_lkp_pSILAC-DIA_btz_39_DIA_118.mzML
Wrote 3785 peptides identified at 1.0% FDR
Read 20200828_QEHFX_lkp_pSILAC-DIA_btz_40_DIA_130.mzML
Wrote 0 peptides identified at 1.0% FDR
Read 20200828_QEHFX_lkp_pSILAC-DIA_btz_40_DIA_157.mzML
Wrote 17870 peptides identified at 1.0% FDR
Read 20200828_QEHFX_lkp_pSILAC-DIA_btz_04_DIA_153.mzML
Wrote 16792 peptides identified at 1.0% FDR
Write Library 20200828_QEHFX_lkp_pSILAC-DIA_btz_QUANT.elib

### QUANTIFY SILAC PAIRS USING SKYLINE

7. Launch Skyline and open a blank document. Ensure that you are in protein mode and default settings (Settings > Default).

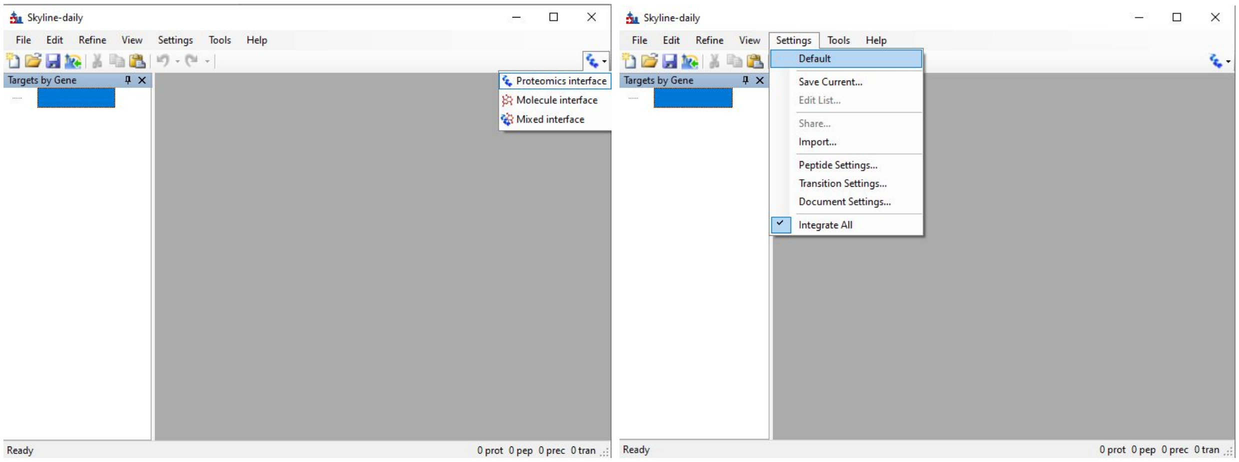


8. Prepare the Skyline document with the appropriate settings per the EncyclopeDIA search performed above and as dictated by the instrument method settings. For this experiment, match the following parameters in Settings > Transition Settings:

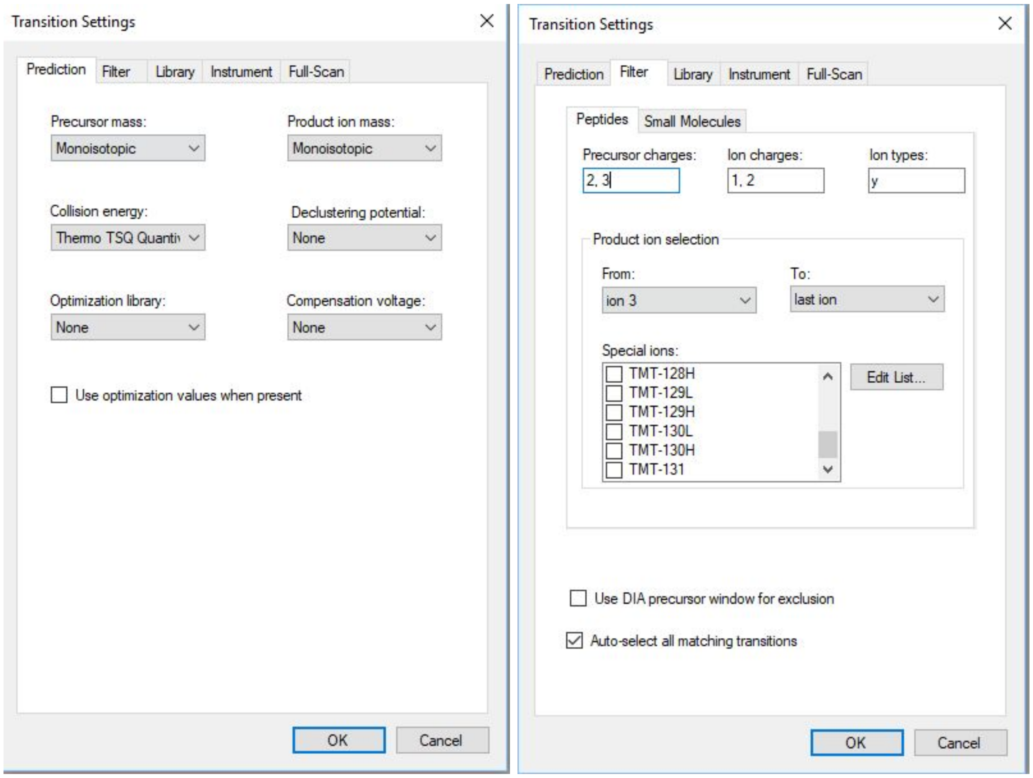

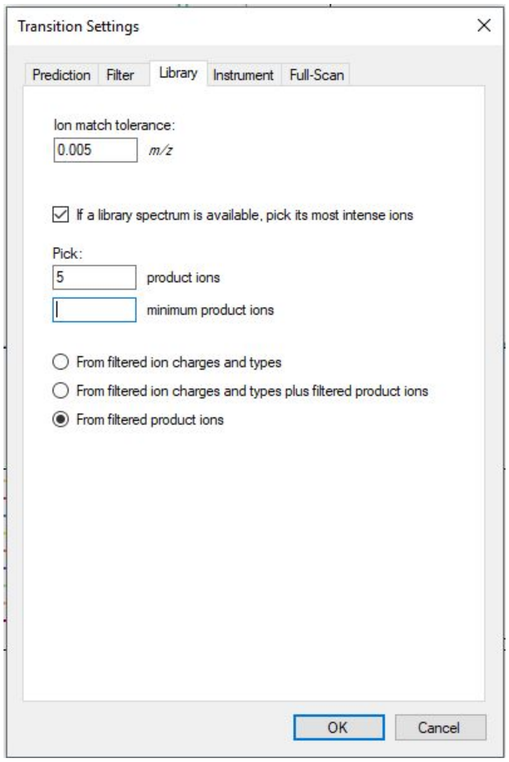


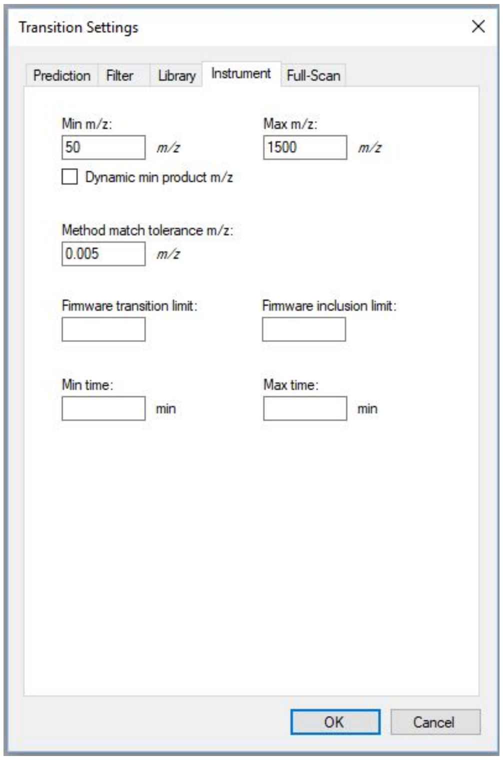


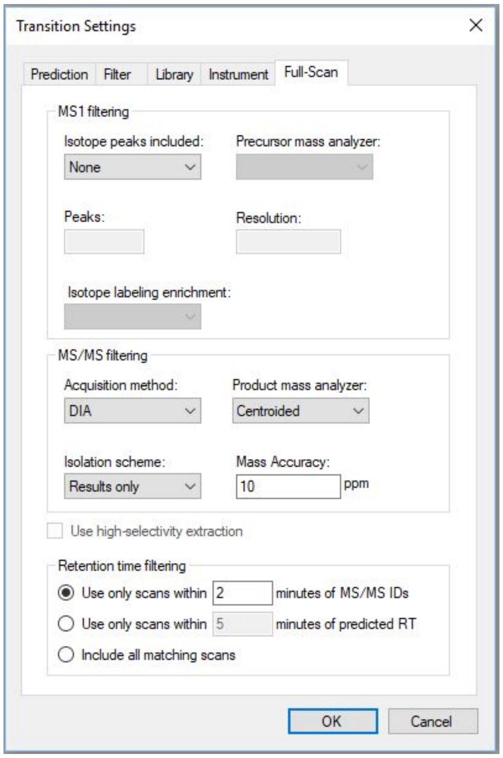


- Prediction: Precursor/Product ion mass=”Monoisotopic”
- Filter: Ion charges=1,2 Ion types=”y”*, From=Ion 3, To=last ion, no special ions

- Library: Ion match tolerance=0.005 m/z, check “If a library spectrum is available, pick its most intense ions”, pick=5 product ions, select “From filtered product ions”
- Instrument: “Min m/z=50, Max m/z=1500, Method match tolerance m/z=0.005
- Full-Scan (MS1): “Isotope peaks included=None”
- Full-Scan (MS/MS): “Acquisition method=DIA, Product mass analyzer=Centroided, Isolation scheme=Results only, Mass Accuracy=10 ppm, check “Use only scans within “2” minutes of MS/MS IDs

9. Next, set the parameters for the Prediction and Filter tabs of the peptide settings (Settings > Peptide Settings), again per the EncyclopeDIA search performed above and as dictated by the instrument method settings. For this experiment:

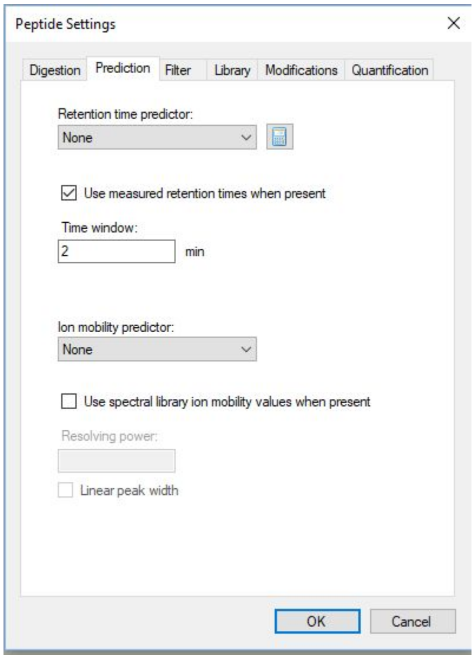


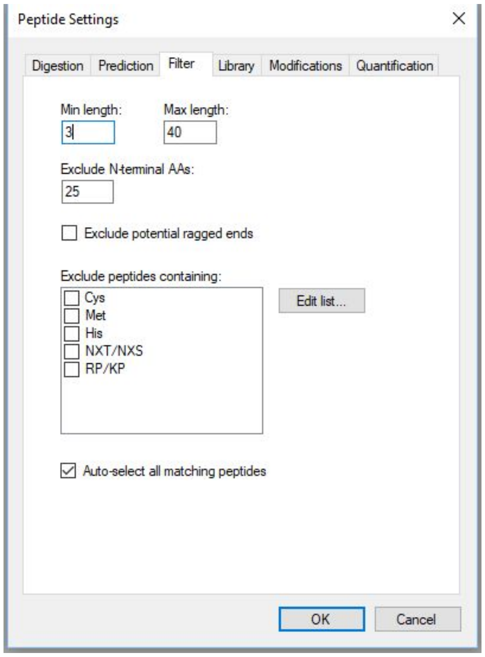


- Prediction: Check “Use measured retention times when present”, Time window=2
- Filter: Min length=3, Max length=40, no excluded amino acids checked

10. In the Digestion tab of the Peptide Settings (Settings > Peptide Settings > Digestion) expand the “Background proteome:” drop down to select <Add…>. Type a working name for this background proteome and then click “Create...” to give the background proteome file a name and filepath location. Click “Add…” and navigate to the same FASTA used in steps 1 and 4 above. Once the background proteome file has been created, press “OK” to return to the Peptide Settings window.

    ***! NOTE****: After selecting the FASTA, this may take some time. Skyline should display a progress bar.*

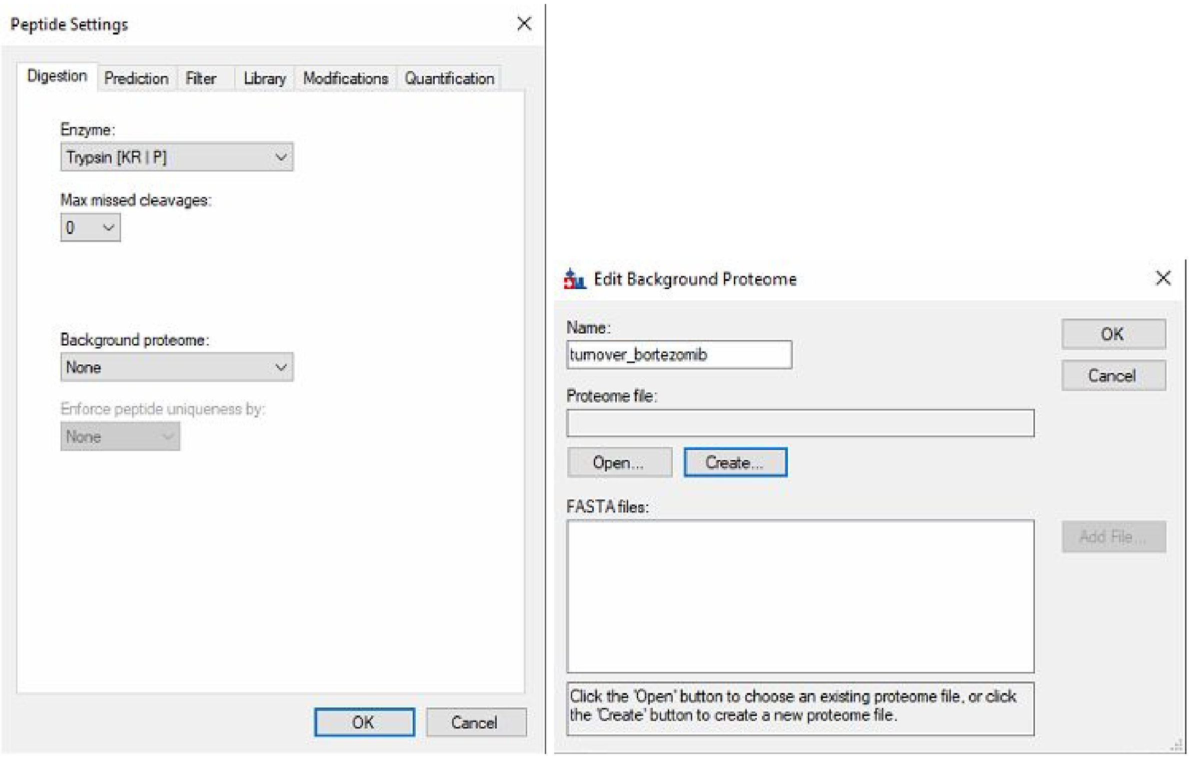


11. In the Library tab of the Peptide Settings window (Settings > Peptide Settings > Library) click “Edit list...” and then “Add…”

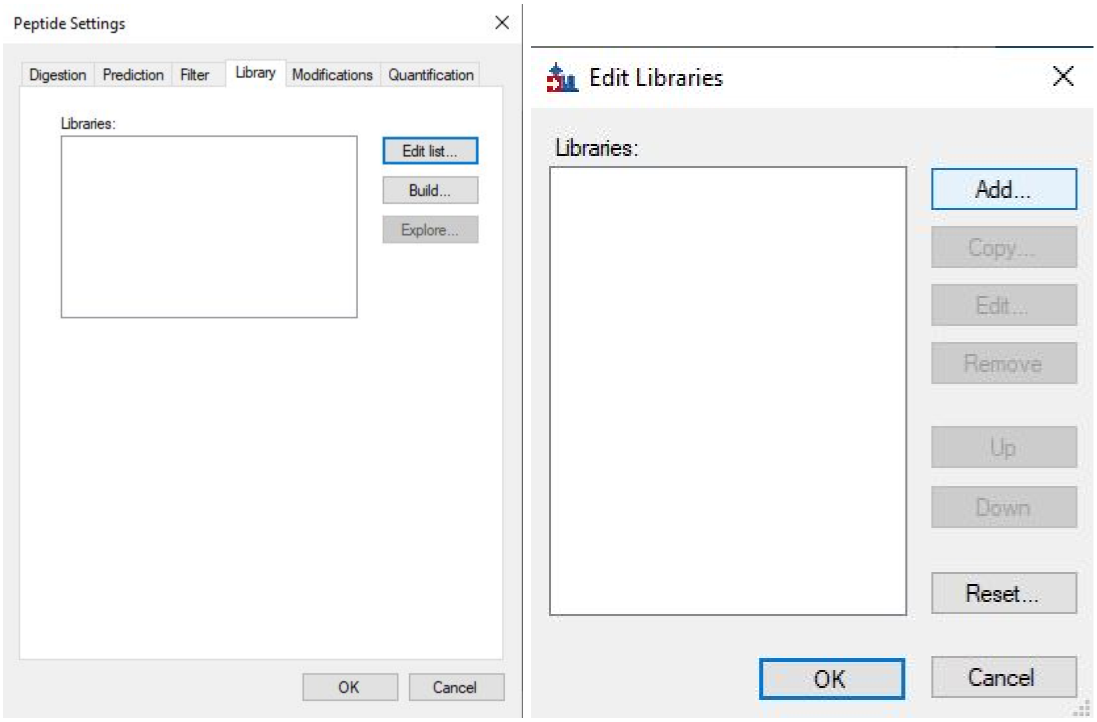

12. Fill out a working name for the library, then next to “Path:” click “Browse…” and navigate to the final ELIB that was saved in step 6 above. Select “OK” and then “OK” again to get back to the Peptide Settings window. Check the box next to the library that was just set up, ensure that the dropdown for “Pick peptides matching:” has “Library” selected, and then click “Explore…”

   ***! NOTE**: After clicking “Explore…” a pop-up may appear notifying that “Peptide settings have been changed. Save changes?” Click YES*

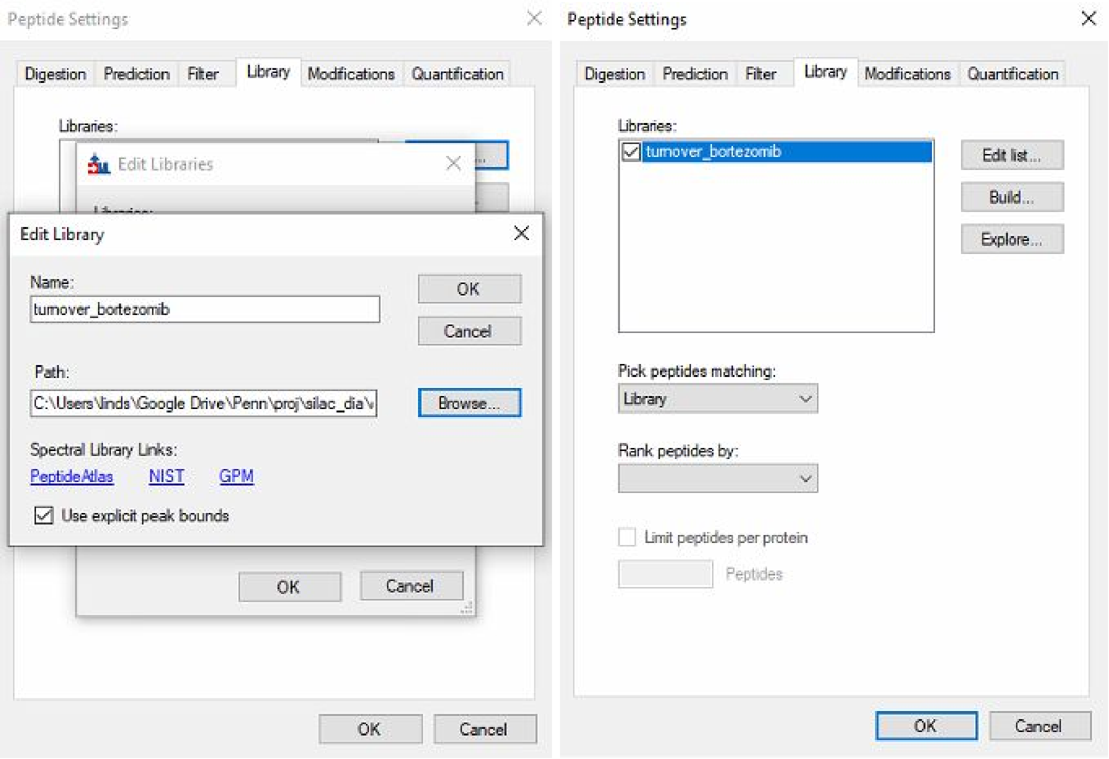


13. Once the spectral library explorer window is launched, check the box at the bottom to “Associate proteins” and then click “Add all…”

   ***! NOTE**: After clicking “Add all…”, a popup progress bar for “Matching peptides to the current document settings” should appear. This may take some time.*

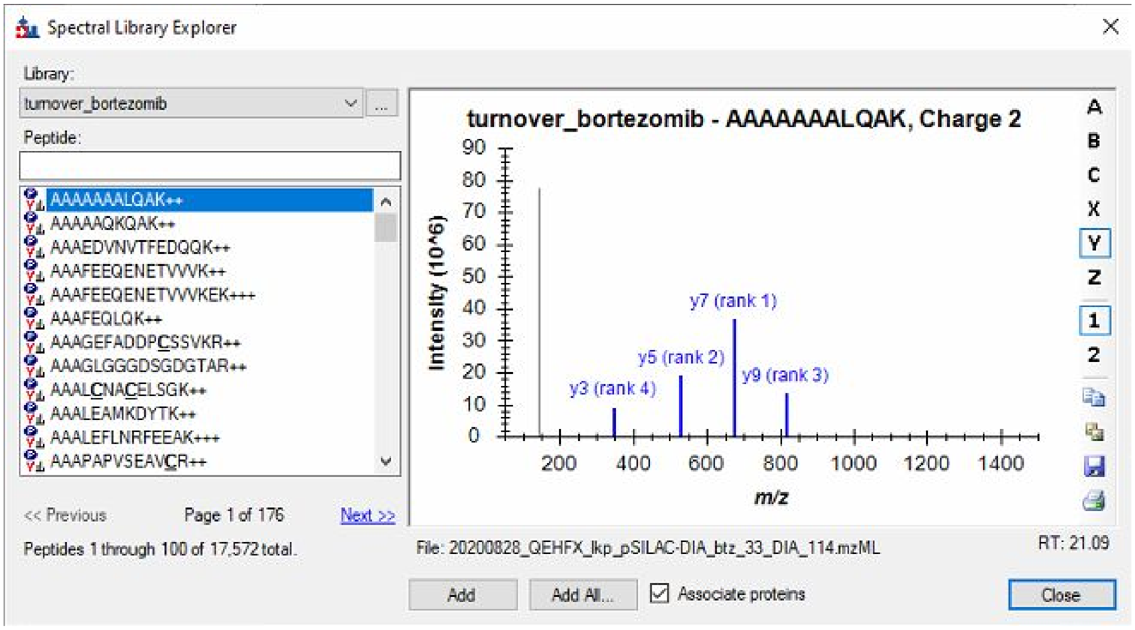


14. When the protein association is complete, a popup window should appear with options for how to handle certain peptide-to-protein mapping situations. Skyline does not use protein groups, so “Add to all matching proteins” is recommended for downstream analysis to ensure that peptide mapping is possible. Click “OK”. After a moment, another popup will appear describing the final results. Select “Add All”. Close out of the spectral library explorer.

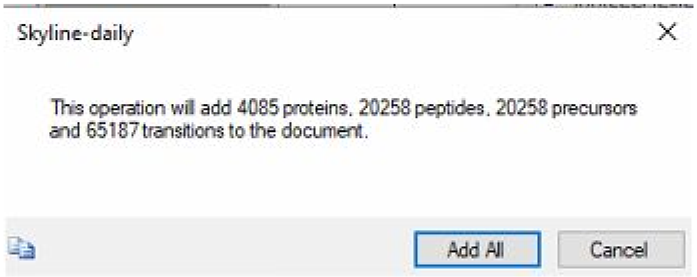


15. The left-hand “Target List” pane should now be populated with proteins, but all these proteins are light/endogenous. Go to Settings > Peptide Settings > Modifications. Under “Isotope modifications”, check the box corresponding to the experiment SILAC labeling scheme, here “heavy” 13C(6)15N(4) arginine and “heavy” 13C(6)15N(20) lysine. Under the “Isotope label type” dropdown, select “heavy”. Click OK to exit out of the Peptide Settings window.

***! NOTE**: Less conventional or custom isotope modifications can be added with Edit list > Add > Edit Isotope Modifications.*

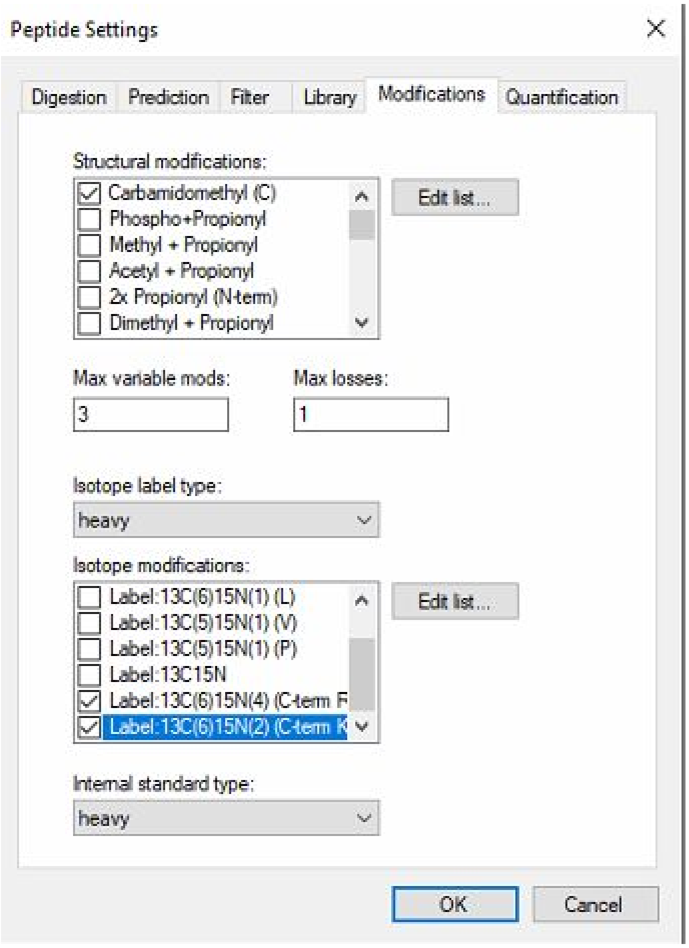


16. Navigate to Refine > Advanced and under the “Remove label type” options, check the box next to “Add” and select “heavy” from the dropdown. Click “OK”.

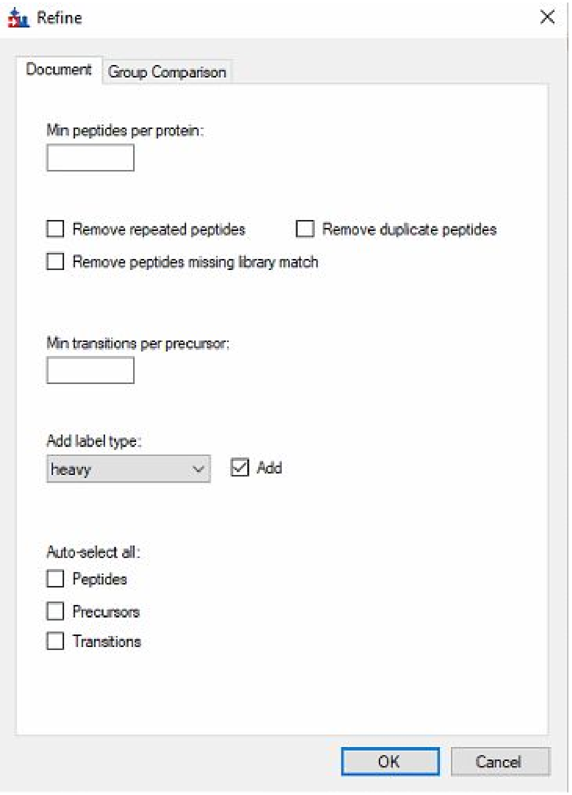


17. Save the Skyline document. Then, choose File > Import > Results to add single-injection replicates and click OK. Navigate to and select all the single-shot, wide-window MZML files used in the EncyclopeDIA analysis above. Choose whether to shorten the file names or not, click “OK”, and the import should begin.

    ***! NOTE**: The chromatogram import graphic should appear and can be used to track progress. This may take time.*

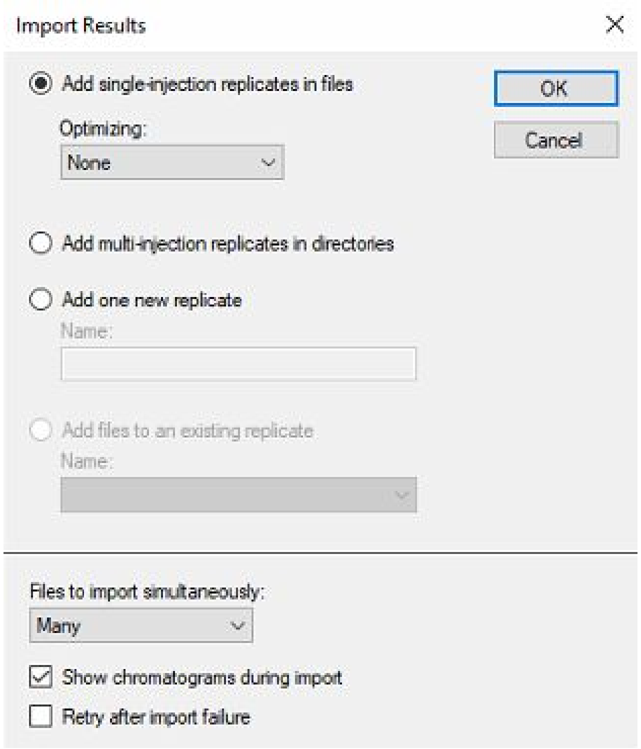


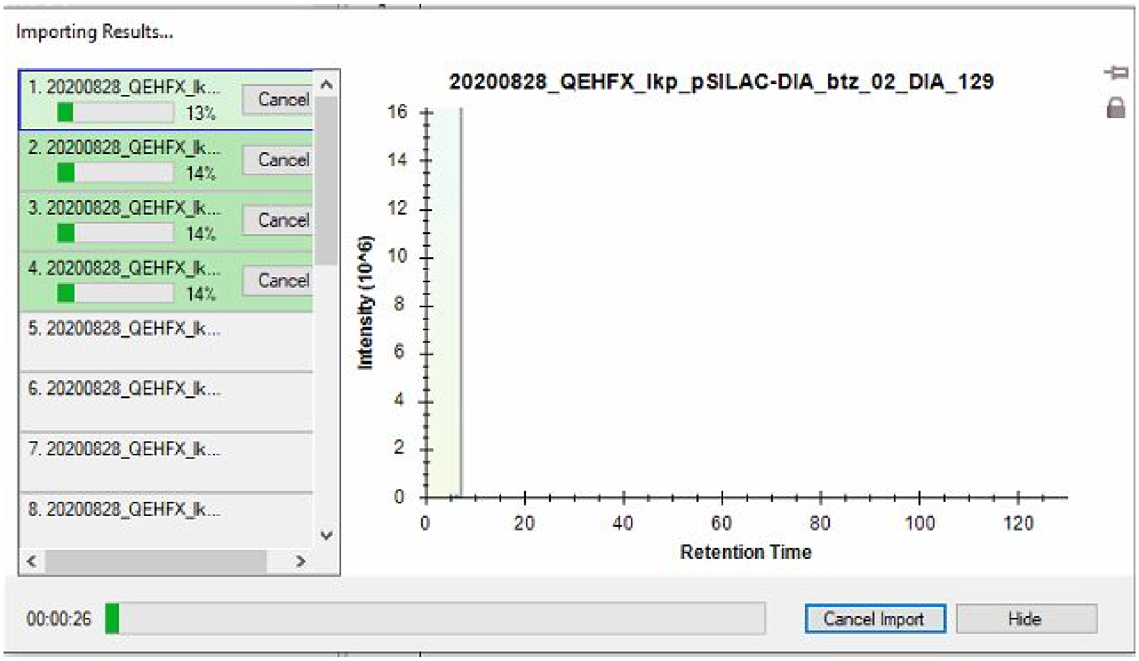


18. Paired SILAC peptides can now be viewed in Skyline. Quantifications can be exported using File > Export > Report...
